# Non-uniform dystrophin re-expression after CRISPR-mediated exon excision in the dystrophin/utrophin double-knockout mouse model of DMD

**DOI:** 10.1101/2022.01.25.477678

**Authors:** Britt Hanson, Sofia Stenler, Nina Ahlskog, Katarzyna Chwalenia, Nenad Svrzikapa, Anna M. L. Coenen-Stass, Marc S. Weinberg, Matthew J. A. Wood, Thomas C. Roberts

## Abstract

Duchenne muscular dystrophy (DMD) is the most prevalent inherited myopathy affecting children, caused by genetic loss of the gene encoding the dystrophin protein. There are currently four FDA-approved drugs for DMD that aim to restore expression of dystrophin by exon skipping using splice switching oligonucleotides. While these therapies require lifelong repeat administration, recent advancements in gene editing technologies have raised the possibility of achieving ‘permanent exon skipping’, and thereby curing the disease with a single treatment. Here we have investigated the use of the *Staphylococcus aureus* CRISPR/Cas9 system and a double-cut strategy, delivered using a pair of AAV9 vectors, for dystrophin restoration in the severely-affected dystrophin/utrophin double knock-out (dKO) mouse. Single guide RNAs were designed to induce double-strand DNA breaks on either side of *Dmd* exon 23, such that the intervening exon 23 sequence is excised when the flanking intronic regions are joined via the non-homologous end joining repair pathway. Exon 23 deletion was confirmed at the DNA level by PCR and Sanger sequencing, and at the RNA level by RT-qPCR. Restoration of dystrophin protein expression was demonstrated by western blot and immunofluorescence staining in mice treated via either intraperitoneal or intravenous routes of delivery. Dystrophin restoration was most effective in the diaphragm, where a maximum of 5.7% of wild-type dystrophin expression was observed. CRISPR treatment was insufficient to extend lifespan in the dKO mouse, and dystrophin was expressed in a within-fiber patchy manner in skeletal muscle tissues. Further analysis revealed a plethora of non-productive DNA repair events, including AAV genome integration at the CRISPR cut sites. This study highlights potential challenges for the successful development of CRISPR therapies in the context of DMD.

## Introduction

Mutations in the gene encoding the dystrophin protein (*DMD*) lead to Duchenne muscular dystrophy (DMD) and Becker muscular dystrophy (BMD). DMD is a lethal, currently incurable, progressive muscle wasting condition and the most common inherited myopathy affecting children^1^. Loss of dystrophin leads to myofiber fragility, cycles of muscle turnover, persistent inflammation, and fibro/fatty muscle degeneration ^2–4^. DMD patients suffer from progressive muscle-wasting and typically become wheelchair-bound around the age of twelve ^5^. While improvements in disease management have extended lifespan, patients typically still succumb to cardiac or respiratory failure around the age of thirty ^6–8^. In contrast, disease severity in BMD patients varies widely, although it is generally considered to be much less severe than DMD, with a later onset of symptoms ^9^. In some rare cases, BMD patients are effectively asymptomatic, despite large internal deletions within the *DMD* gene ^10–12^. The molecular reason for the differences in pathological severity between these dystrophinopathies relates to the nature of their respective causative mutations ^13^. In DMD, the dystrophin translation reading frame is typically disrupted, often by whole exon deletions. In the case of BMD, the dystrophin mutation often does not affect the translation reading frame, and so some degree of dystrophin functionality is retained ^14^.

The *DMD* gene consists of 79 exons, many of which encode for redundant structural domains which are not essential for dystrophin function ^15^. DMD is therefore amenable to splice correction therapy in which short antisense oligonucleotides targeting splicing motifs in the dystrophin pre-mRNA effectively mask specific exons from the splicing machinery, leading to their exclusion from the processed mRNA (i.e. exon skipping) ^16^. This alternative splicing event restores the translation reading frame and rescues expression of a truncated, and partially functional, pseudo-dystrophin. The aim of exon skipping is therefore to convert the DMD phenotype to a milder BMD-like phenotype. To date, there are four FDA-approved exon skipping drugs (eteplirsen, golodirsen, viltolarsen, and casimersen) ^17^. These therapies are currently very expensive (∼$300,000 per patient *per annum*), show limited efficacy (i.e. restoration of ∼1% of normal dystrophin protein levels for eteplirsen ^18^), and require lifelong administration. There is therefore intense interest in more permanent dystrophin restoration strategies, such as gene editing.

In recent years, the gene editing field has experienced a revolution with the discovery and development of the CRISPR/Cas9 (clustered regularly interspaced short palindromic repeats/CRISPR associated protein 9) system ^19–21^, for which the Nobel Prize in Chemistry was awarded to Doudna and Charpentier in 2020. In the most commonly-used configuration, the CRISPR approach requires two components: (i) the Cas9 enzyme which induces double-strand breaks (DSBs) in genomic DNA, and (ii) a single guide RNA (sgRNA) that specifies the target DNA sequence. The use of an RNA component makes this technology highly versatile as there are relatively few constraints on what sequence can be targeted, and sgRNAs can be generated rapidly. Furthermore, the CRISPR/Cas9 system enables facile multiplexing of genome editing events, such that multiple loci can be targeted simultaneously ^20^. CRISPR-induced DSB lesions are resolved by cellular DNA damage repair pathways, the most common being the error-prone non-homologous end joining (NHEJ) pathway, and homology directed repair (HDR). CRISPR-based approaches for DMD have mainly focused on strategies relying on the NHEJ DNA repair pathway, as HDR is generally not active in quiescent or terminally differentiated cells ^22^. The leading approaches are (i) a double-cut strategy in which a target exon(s) is excised, essentially achieving ‘permanent exon skipping’ ^23–28^, and (ii) a single-cut strategy aimed at inducing an indel for the purposes of either disrupting a splice signal, or reframing an exon ^29–34^. *In vivo* delivery of the Cas9 and sgRNA transgenes is typically achieved with the use of adeno-associated viral (AAV) vectors ^35^, of which there are several muscle-tropic serotypes ^36, 37^. AAVs have been used extensively for gene therapy applications and have already demonstrated safe and efficacious delivery in clinical trials ^38, 39^. Notably, the commonly used *S. pyogenes* Cas9 (SpCas9) protein is too large to be effectively packaged into AAV vectors, which has led to the development of smaller orthogonal Cas9 proteins^40^. The compact (3.2 kb) Cas9 derived from *Staphylococcus aureus* (i.e. SaCas9) has emerged as the most widely-utilized variant of choice ^41^.

The most commonly-used mouse model of DMD is the *mdx* mouse, which recapitulates some aspects of DMD pathology. Specifically, a ‘crisis’ period of extensive myonecrosis and muscle regeneration is observed between 2 and 5 weeks of age ^42^, with more severe histopathological features reminiscent of DMD pathology (i.e. fibro/fatty degeneration) observed much later (∼80 weeks) ^43^. Cardiomyopathy is generally not observed in the *mdx* mouse unless experimentally induced ^44^. Importantly, the *mdx* mouse exhibits only a small decrease in lifespan, and impairment of muscle function is relatively mild. Disease in the *mdx* mouse is similar to that observed in other dystrophin-null models ^45^. In contrast, the dKO mouse (a double knock-out of dystrophin and its paralog utrophin) exhibits much more severe pathology, including kyphosis, respiratory difficulties, and premature death (animals typically die ∼8-10 weeks of age) ^46–49^. Notably, exon skipping therapy to rescue dystrophin protein expression in the dKO mouse reverses dystrophic pathology and dramatically extends lifespan ^48, 49^. Importantly, the dKO mouse carries an identical genetic mutation to that of the *mdx* mouse (i.e. a nonsense mutation in *Dmd* exon 23) and so these two strains can be treated using common exon skipping or CRISPR-based therapeutic strategies. Here, we report *Dmd* gene editing and dystrophin restoration in the severely-affected dKO mouse model of DMD using an AAV-delivered (wo vector), double-cut CRISPR strategy. However, the observed levels of gene editing were insufficient to extend lifespan in the dKO mouse, or to correct circulating miRNA biomarker levels. Dystrophin protein expression in the treated animals was localized to the sarcolemma, but exhibited a within-fiber patchy distribution pattern. Long-read sequencing of the editing target site revealed a plethora of non-productive editing outcomes, including AAV backbone integration events and indels which disrupt the sgRNA target sites. We propose that inefficient editing using this approach will inevitably lead to the generation of chimeric fibers containing both dystrophin-expressing and non-dystrophin-expressing myonuclei, and that the resulting incomplete pattern of sarcolemmal dystrophin may be insufficient to correct the dystrophic phenotype.

## Results

### Screening of *Dmd*-targeting guide RNAs in murine fibroblasts

In order to correct the genetic defect in the dKO mouse we have utilized a multiplex genome editing strategy intended to excise *Dmd* exon 23, similar to approaches described previously ^23– 25, 27, 31^. This approach aims to induce DNA DSBs on either side of the target exon with the resulting genetic lesion repaired via the NHEJ pathway. The expected productive editing outcome is the excision of exon 23 and the fusion of the flanking introns (**Figure 1A**). Putative SaCas9 PAM sites (i.e. NNGRRT ^41^, where R = A or G) were identified in the intronic sequence surrounding the exon 23. Four potential target sites were selected (two each on either side of the exon) and sgRNAs cloned into U6 expression cassettes (**Figure 1B**). To identify the optimal dual guide combination, mouse fibroblast cells (10T1/2) were transfected with SaCas9 and all four possible combinations of the sgRNA plasmids (**Figure 1C**). Genome editing was assessed by PCR using primers which span the edit site and produce an 833 bp amplicon. Excision of *Dmd* exon 23 was observed for two of the four combinations, as indicated by the detection of a smaller PCR product ∼446 bp. The optimal combination of sgRNAs (2+4) exhibited gene editing efficiency of ∼4.7%, and was selected for further development. The edited amplicon was analyzed by direct Sanger sequencing to characterize NHEJ-induced indels (**Figure 1D**). The major product in the sequencing chromatogram corresponded to seamless repair of the up- and downstream introns at the expected sgRNA cut sites (i.e. with no indel), although an increase in background signal was observed around the repair site indicative of low level of indel formation.

**Figure 1.**
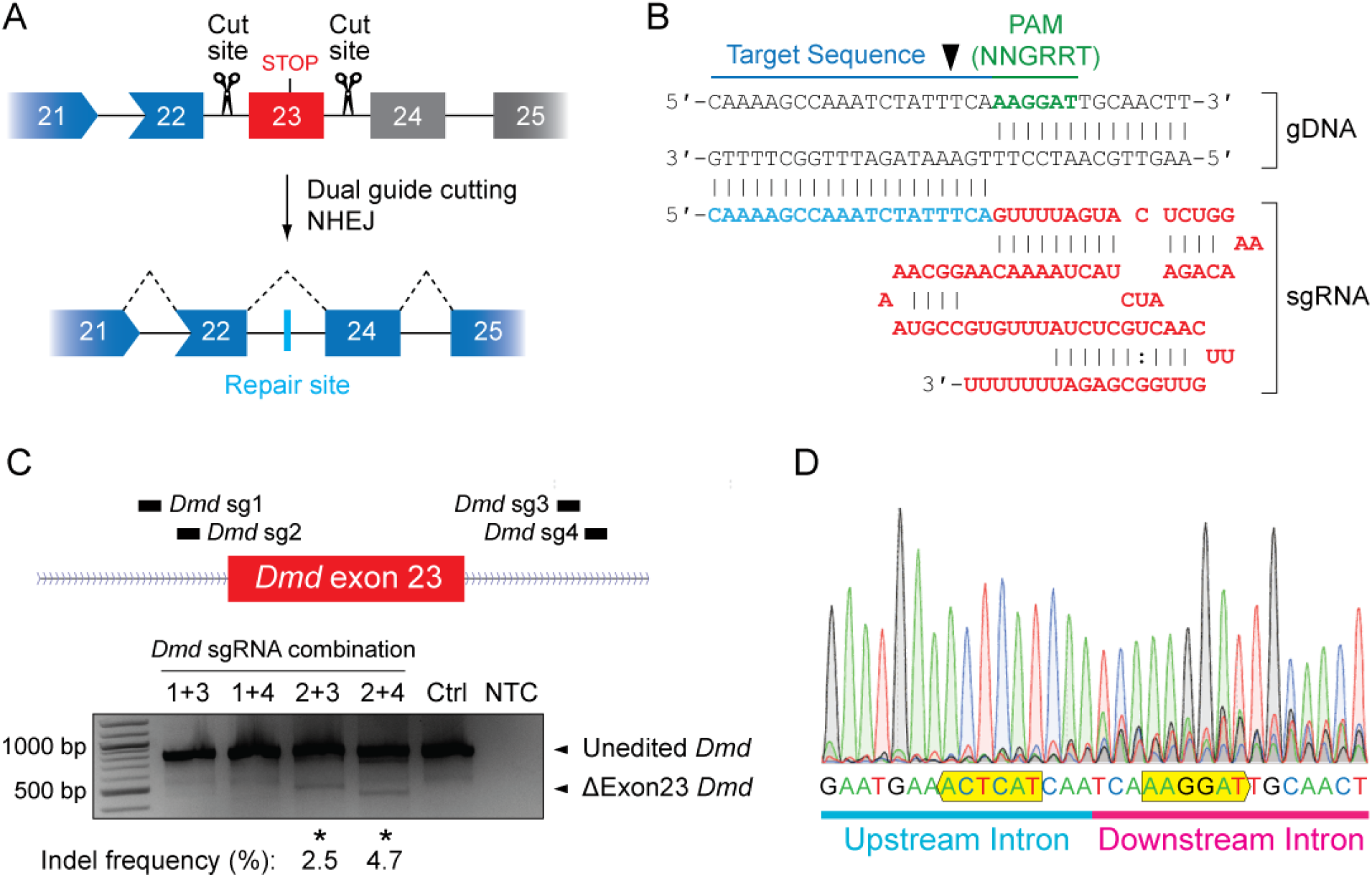
CRISPR/Cas9-mediated *Dmd* exon 23 excision in cell culture. (**A**) Schematic of dual sgRNA CRISPR/Cas9-mediated excision of murine *Dmd* disease-causing exon 23 (in red), resulting in NHEJ and a repair indel within the intron that restores the *Dmd* translation reading frame. Out-of-frame exons are indicated in grey. (**B**) Schematic of the structure of an SaCas9 sgRNA in complex with its cognate gDNA target site. The SaCas9-specific protospacer adjacent motif (PAM) sequence (NNGRRT, where R is A or G) is shown in green and is located on the non-targeted DNA strand. (**C**) Screening of the activity of four combinations of sgRNAs for *Dmd* exon 23 excision in murine 10T1/2 fibroblast cells. The genomic target site was amplified by PCR and the products resolved by agarose gel electrophoresis. Successful editing is indicated by a shorter, ΔExon23 *Dmd* product and editing percentages indicated for successful sgRNA combinations. (**D**) Sanger sequencing chromatogram confirming successful exon 23 excision and fusion of the up- and downstream intronic regions following treatment with SaCas9 sgRNA combination 2+4.

### CRISPR-mediated dystrophin rescue in the dKO mouse after intraperitoneal injection

To evaluate the potential for CRISPR/Cas9-mediated *Dmd* editing in the severely-affected dKO (double knockout for dystrophin and utrophin) mouse *in vivo*, two separate AAV9 vectors were designed. The first vector encodes for an SaCas9 transgene with compact regulatory sequences to minimize the size of the AAV genome (i.e. the CMV promoter and SV40 minimal poly(A) signal). The total size of the resulting AAV genome (including ITRs) was 4,806 bp, which is below the maximum limit for AAV packaging (estimated at 5.2 kb) ^50^. For the second AAV vector, two different RNA polymerase III promoters (the human H1 and U6 snRNA promoters) were selected to drive expression of the *Dmd*-targeting sgRNAs in order to minimize the repetitive sequence within the AAV genome (**Figure 2A**).

**Figure 2.**
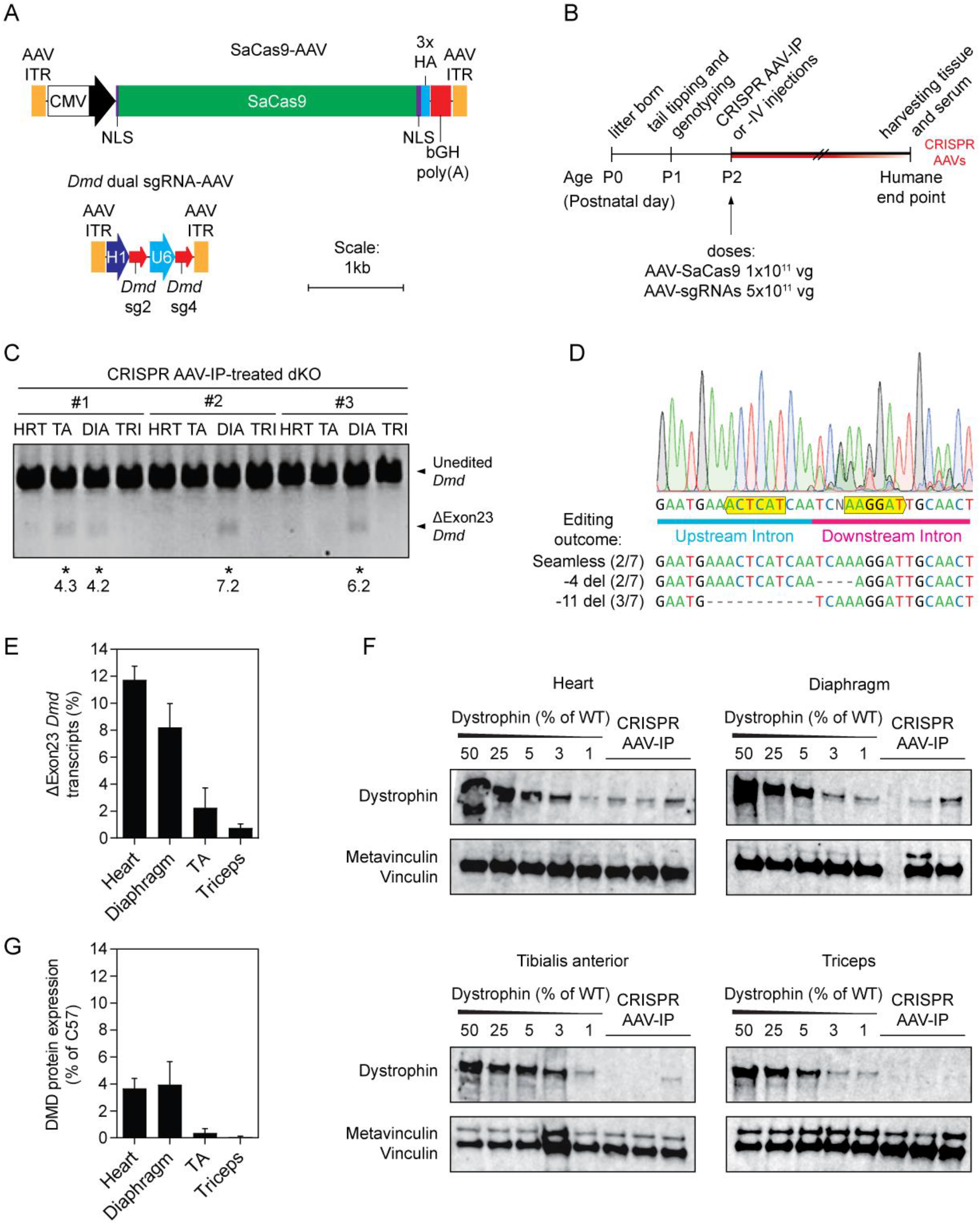
Evaluation of *Dmd* exon 23 excision and dystrophin restoration in dKO mice after intraperitoneal injection of neonates. (**A**) Schematics of the SaCas9-AAV and *Dmd* dual sgRNA-AAV genomes contained within AAV2 inverted terminal repeats (ITRs). The SaCas9 gene is driven by a CMV promoter and enhancer, is flanked by an SV40 nuclear localization signal (NLS) on the N-terminus and a nucleoplasmin NLS on the C-terminus, is tagged with three HA tags and is terminated with a bGH poly(A) signal. Expression of *Dmd* sg2 and sg4 is driven by an H1 and a U6 promoter, respectively (sgRNAs selected based on results in Figure 1). (**B**) Treatment timeline for dKO mice with CRISPR AAV9 particles. Mice were treated by intraperitoneal (IP) injection (*n*=3) at postnatal day two (P2) with 1×10^11^ vg of AAV-SaCas9 and 5×10^11^ vg of AAV-sgRNA and sacrificed at the humane end point. DNA, RNA, and protein were extracted from the heart (HRT), diaphragm (DIA), tibialis anterior (TA), and triceps (TRI) muscles. (**C**) Visualization of successful CRISPR-mediated *Dmd* exon 23 excision (ΔExon23 *Dmd*) in the gDNA by PCR and agarose gel electrophoresis. Samples exhibiting the desired editing outcome are highlighted with asterisks and corresponding indel frequencies indicated. (**D**) A representative Sanger sequencing chromatogram confirming successful exon 23 excision and fusion of the up- and downstream intronic regions following CRISPR AAV treatment. PAM sites are labelled with yellow boxes. (**E**) Reverse transcriptase-quantitative PCR (RT-qPCR) indicating the levels of ΔExon23 *Dmd* transcripts relative to WT. Western blots (**F**) and quantification of dystrophin protein restoration (**G**) relative to the vinculin protein loading control. Values are mean+SEM.

Postnatal day 2 (P2) dKO mice were treated with 1×10^11^ vector genomes (vg) of SaCas9-AAV and 5×10^11^ vg of dual sgRNA-AAV, by intraperitoneal (IP) injection (**Figure 2B**). Treated mice were monitored until they reached the humane end-point, at which time the heart, diaphragm, tibialis anterior (TA), and triceps were harvested in order to determine the efficacy of gene editing at the gDNA, RNA, and protein levels.

The ΔExon23 *Dmd* gene product, indicative of productive editing at the genomic target site, was detected sporadically across the tissues analyzed by PCR (**Figure 2C**), and successful excision and repair of the flanking intronic regions at the cut sites was confirmed by Sanger sequencing of the bulk edited PCR amplicon (**Figure 2D**). The sequencing chromatogram demonstrated seamless fusion of the flanking intronic regions, with an increase in background signal around the repair site indicative of low levels of indel formation. Cloning of the edited amplicon and sequencing of seven individual clones revealed three major editing events: seamless repair (2/7), a 4 bp deletion downstream of the fusion site (2/7), and an 11 bp deletion upstream of the fusion site (3/7) (**Figure 2D**).

Successful excision of the gDNA encoding *Dmd* exon 23 is expected to manifest as permanent ‘exon skipping.’ As such, levels of ΔExon23 *Dmd* mRNA transcripts were determined by RT-qPCR using primers spanning exons 22-24 (i.e. exon ‘skipped’) and spanning exons 23-24 (i.e. exon ‘unskipped’). Successful exon excision was detected in all tissues tested, thereby indicating that this method is more sensitive for detecting editing than end-point PCR analysis of gDNA. ΔExon23 *Dmd* transcript levels were highest in the heart and diaphragm at ∼12% and 8% respectively (**Figure 2E**). Editing in the TA and triceps was less successful, with only ∼2% and 1% ΔExon23 *Dmd* transcripts detected respectively.

A similar pattern was observed at the protein level, whereby dystrophin protein expression was observed in the heart, diaphragm, and TA muscles of the treated dKO mice, but at negligible levels in the triceps as determined by western blot (**Figure 2F,G**). The size of the restored dystrophin protein was similar to that observed in WT controls, which is to be expected as deletion of exon 23 results in a small (71 amino acids, 420 Da) change in the size of the dystrophin protein. Dystrophin was undetectable in saline-treated dKO control animals (**Figure S1**).

Immunofluorescence (IF) staining for dystrophin protein was performed to visualize of the pattern of successful gene editing within CRISPR AAV-treated muscle tissue sections. Transverse sections obtained from each of the four muscle tissues were stained with anti-dystrophin fluorescent antibodies to assess the distribution of restored fibers throughout the selected tissues. Sections were co-stained for laminin (LAMA2), a basal lamina protein that delineates the boundaries of individual myofibers (**Figure 3**). Tissue sections from WT and saline-treated dKO mice were included as positive and negative controls, respectively. Expression of re-expressed dystrophin, localized at the sarcolemma, was confirmed in all four tissues analyzed following CRISPR AAV treatment via IP injection, whereas dystrophin staining was largely absent in sections from all the muscle tissues of saline-treated dKO mice. In the heart and diaphragm, where the highest levels of editing were observed by RT-qPCR and western blot (**Figure 2E-G**), the restored fibers were distributed throughout the tissue section. In the two limb muscles, however, the corrected fibers formed a more clustered pattern, with patches of dystrophin-positive and - negative fibers. This between-fiber ‘patchiness’ may be due to mosaicism of successful gene editing early in the muscle development of these mice, or as a consequence of differences in vascular permeability throughout the muscle tissue. Regions of dystrophin-negative staining within CRISPR-corrected dystrophin-positive myofibers were also observed (**Figure 3**, arrowheads), which was most apparent in the TA and triceps muscles. Notably, we observed IF signal even when dystrophin levels were very low as determined by western blot, consistent with the former technique having higher sensitivity, as has been suggested previously ^51^.

**Figure 3.**
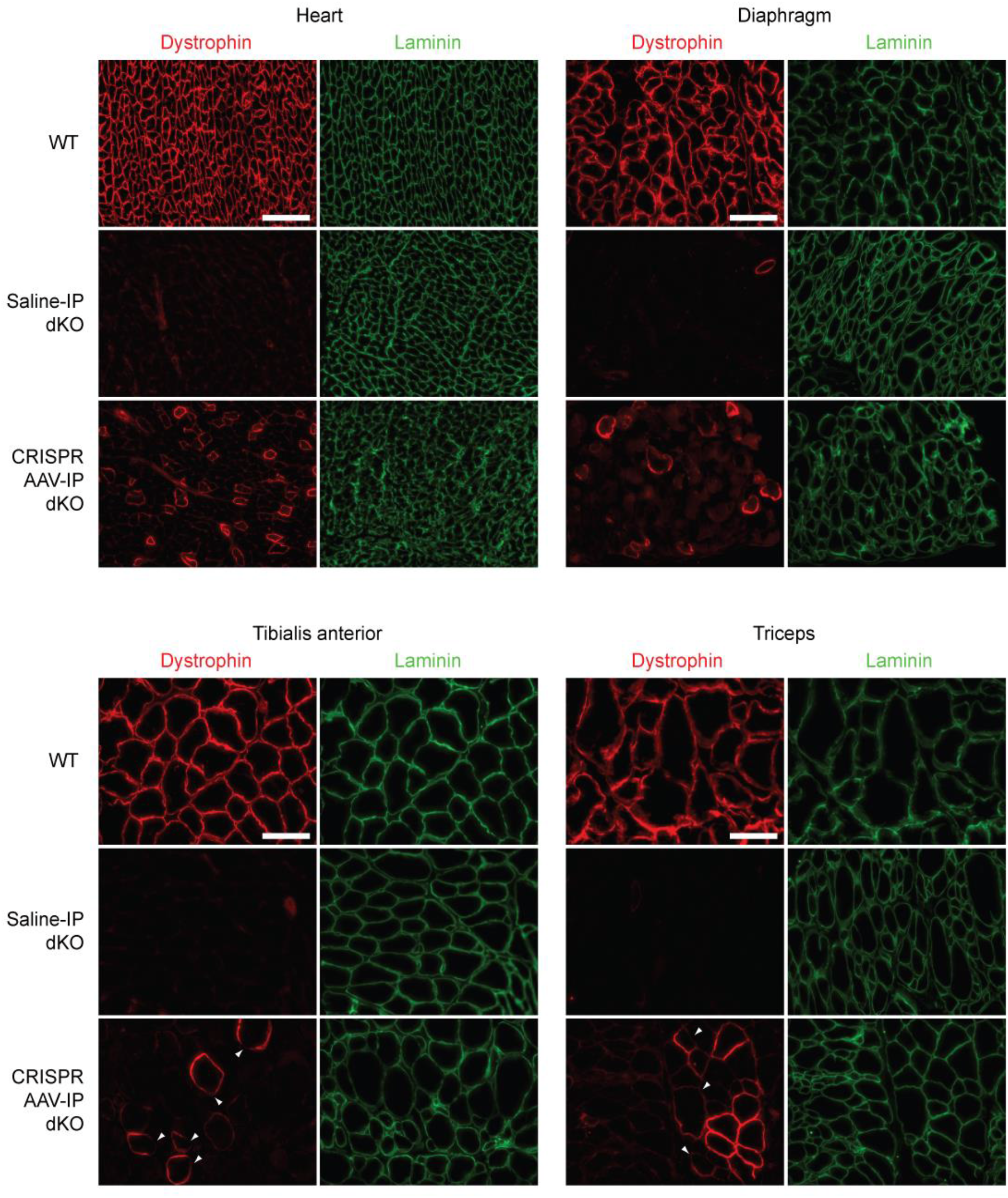
Dystrophin restoration in tissue sections from intraperitoneal CRISPR AAV-treated dKO muscle. Representative dystrophin IF staining of transverse tissue sections obtained from heart, diaphragm, TA, and triceps muscles of dKO mice treated with CRISPR AAV9 particles by IP injection at P2. Untreated wild-type (WT) and saline-treated dKO samples were analyzed in parallel as controls. Sections were co-strained for laminin to delineate myofiber boundaries. Images were taken at 20× magnification. Scale bars represent 100 μm. Arrowheads indicate regions of patchy sarcolemmal dystrophin staining.

### CRISPR-mediated dystrophin rescue in the dKO mouse after intravenous injection

The utility of systemic (intravenous, IV) administration of the CRISPR AAVs was assessed in the dKO mouse model to determine whether broader targeting of muscle tissues could be achieved, and because this is a more widely used method for drug therapy delivery in patients. Several other studies have employed this route of administration for gene editing therapy in neonatal mice, demonstrating the utility of early intervention CRISPR AAV treatment ^23, 24, 27^. An equivalent dose to that used for IP administration of CRISPR AAVs was delivered to P2 dKO mice via the facial vein (**Figure 4B**). Successful excision of *Dmd* exon 23 was detected by PCR in ∼81% (i.e. 13/16) of the tissues analyzed (**Figure 4A**), and successful gene editing was confirmed by Sanger sequencing of the bulk edited PCR product (**Figure 4B**). Cloning of the edited amplicon and sequencing of five individual clones revealed both seamless editing and the formation of various random indels (**Figure 4B**), as is typical of repair by NHEJ.

**Figure 4.**
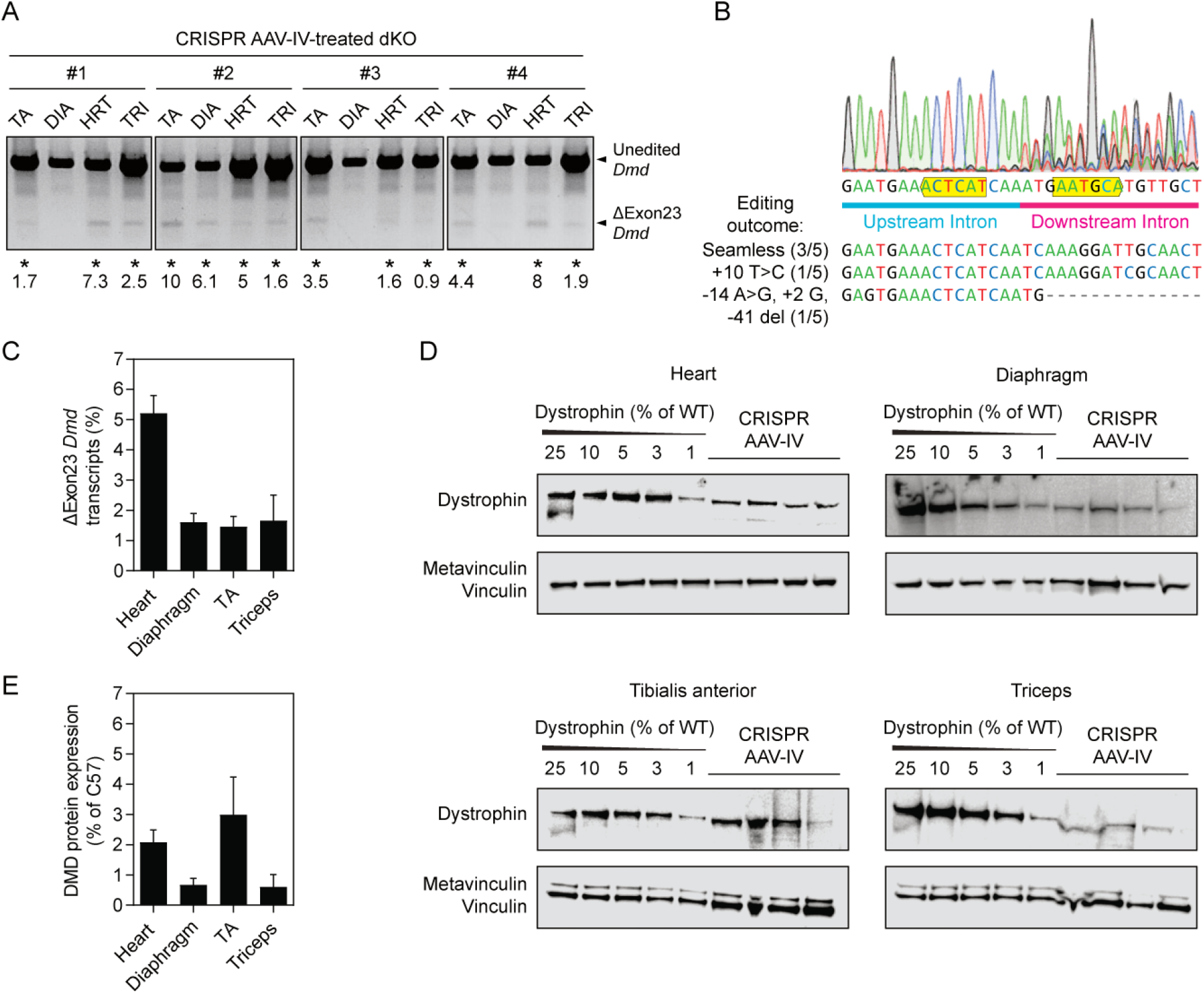
Evaluation of *Dmd* exon 23 excision and dystrophin restoration in dKO mice after intravenous injection of neonates. dKO mice were treated by intravenous (IV) injection (*n*=4) via the facial vein at postnatal day two (P2) with 1×10^11^ vector genomes (vg) of SaCas9-AAV and 5×10^11^ vg of dual sgRNA-AAV and sacrificed at the humane end point. DNA, RNA, and protein were extracted from the heart (HRT), diaphragm (DIA), tibialis anterior (TA), and triceps (TRI) muscles. (**A**) Visualization of successful CRISPR-mediated *Dmd* exon 23 excision (ΔExon23 *Dmd*) in the gDNA by PCR and agarose gel electrophoresis. Samples exhibiting the desired editing outcome are highlighted with asterisks and corresponding indel frequencies indicated. (**B**) A representative Sanger sequencing chromatogram confirming successful exon 23 excision and fusion of the up- and downstream intronic regions following CRISPR AAV treatment. PAM sites are labelled with yellow boxes. (**C**) Reverse transcriptase-quantitative PCR (RT-qPCR) indicating the levels of ΔExon23 *Dmd* transcripts relative to WT. Western blots (**D**) and quantification of dystrophin protein restoration (**E**) relative to the vinculin protein loading control. Values are mean+SEM.

The proportion of ΔExon23 *Dmd* mRNA transcripts was highest in the heart tissue at ∼5%, following IV administration of CRISPR AAVs (**Figure 4C**). Transcripts lacking *Dmd* exon 23 in the diaphragm, TA and triceps were detected at a much lower abundance of ∼1.5% WT levels. Dystrophin protein, as measured by western blot, was partially restored in all four tissues (**Figure 4D,E**) with the highest expression in the heart and TA muscles (∼2% of WT levels) and lowest in the diaphragm and triceps (<1% of WT levels).

IF staining for dystrophin and laminin in transverse sections confirmed that dystrophin was partially restored in the heart, diaphragm, TA, and triceps muscles of neonatal dKO mice treated systemically with CRISPR AAVs by IV injection (**Figure 5**). The presence of dystrophin-re-expressing myofibers was interspersed throughout the muscle in the heart, and to a lesser extent also the diaphragm, while in the TA and triceps there was distinct between-fiber patchiness (i.e. the sections consisted of both dystrophin-positive and dystrophin-negative myofibers). Within-fiber patchiness was also observed, whereby some dystrophin-positive myofibers exhibited incomplete dystrophin staining around the fiber circumference (where laminin co-staining was clearly present) (**Figure 5**, arrowheads).

**Figure 5.**
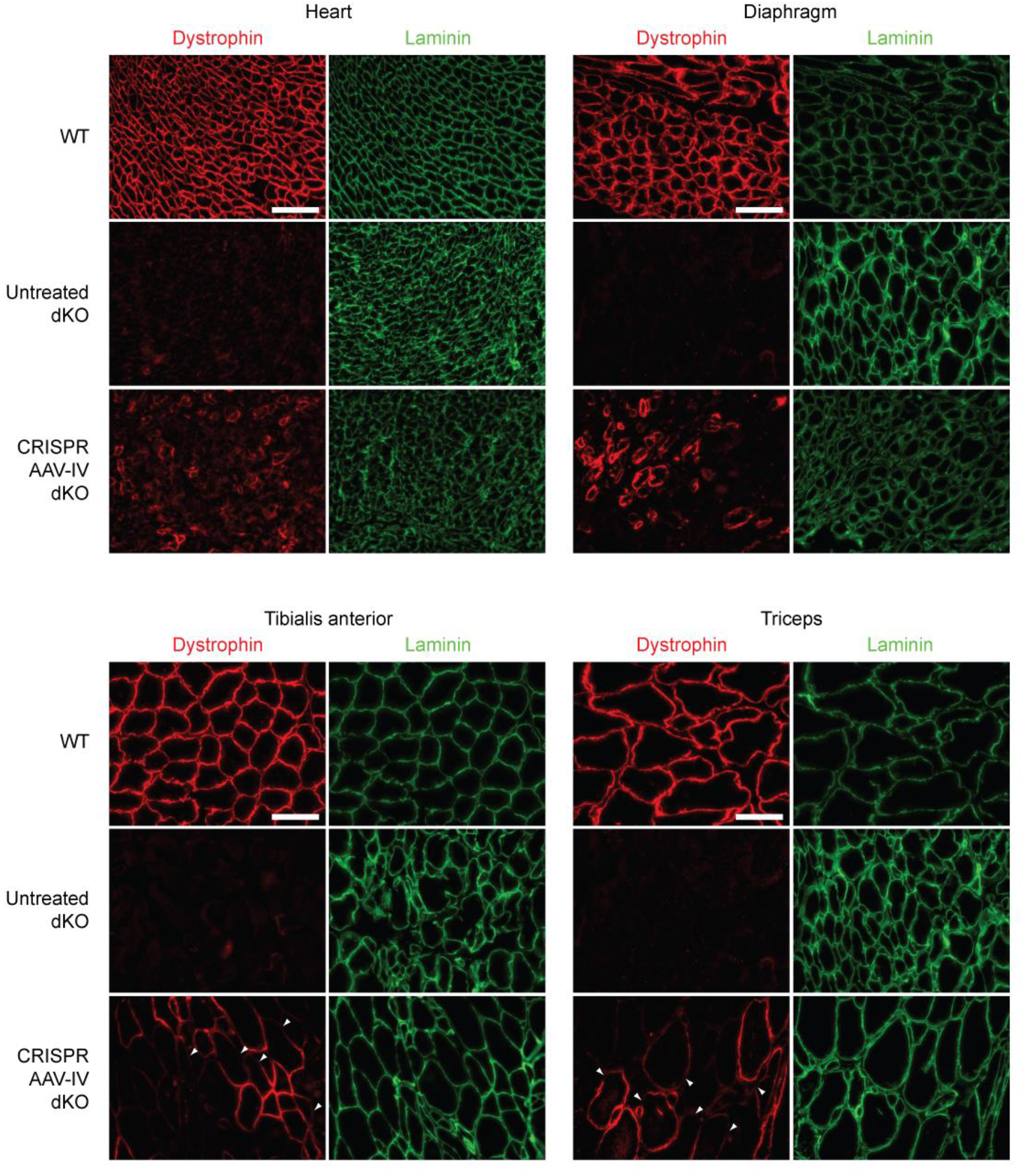
Dystrophin restoration in tissue sections from intravenous CRISPR AAV-treated dKO muscle. Representative dystrophin IF images of transverse tissue sections obtained from heart, diaphragm, TA, and triceps muscles of dKO mice treated with CRISPR AAV9 particles by IV injection at P2. Untreated wild-type (WT) and saline-treated dKO samples were analyzed in parallel as controls. Sections were co-strained for laminin to delineate myofiber boundaries. Images were taken at 20× magnification. Scale bars represent 100 μm. Arrowheads indicate regions of patchy sarcolemmal dystrophin staining.

### Dystrophin gene editing did not improve dKO lifespan

No difference in the weight gain of the CRISPR AAV-treated dKO mice was observed (**Figure 6A**). Furthermore, the average lifespan of the AAV-IP and -IV treatment groups was ∼7-weeks and ∼8-weeks, respectively (**Figure 6B**). This was not significantly different to that of the control dKO mice which also succumbed to the dystrophic pathology around the age of 8-weeks (*P* = 0.7857 and >0.9999 for the AAV-IP and AAV-IV treatment groups relative to the saline control, respectively). Similarly, serum miRNA biomarkers were not restored to WT levels in the CRISPR-treated animals (**Figure S2**).

**Figure 6.**
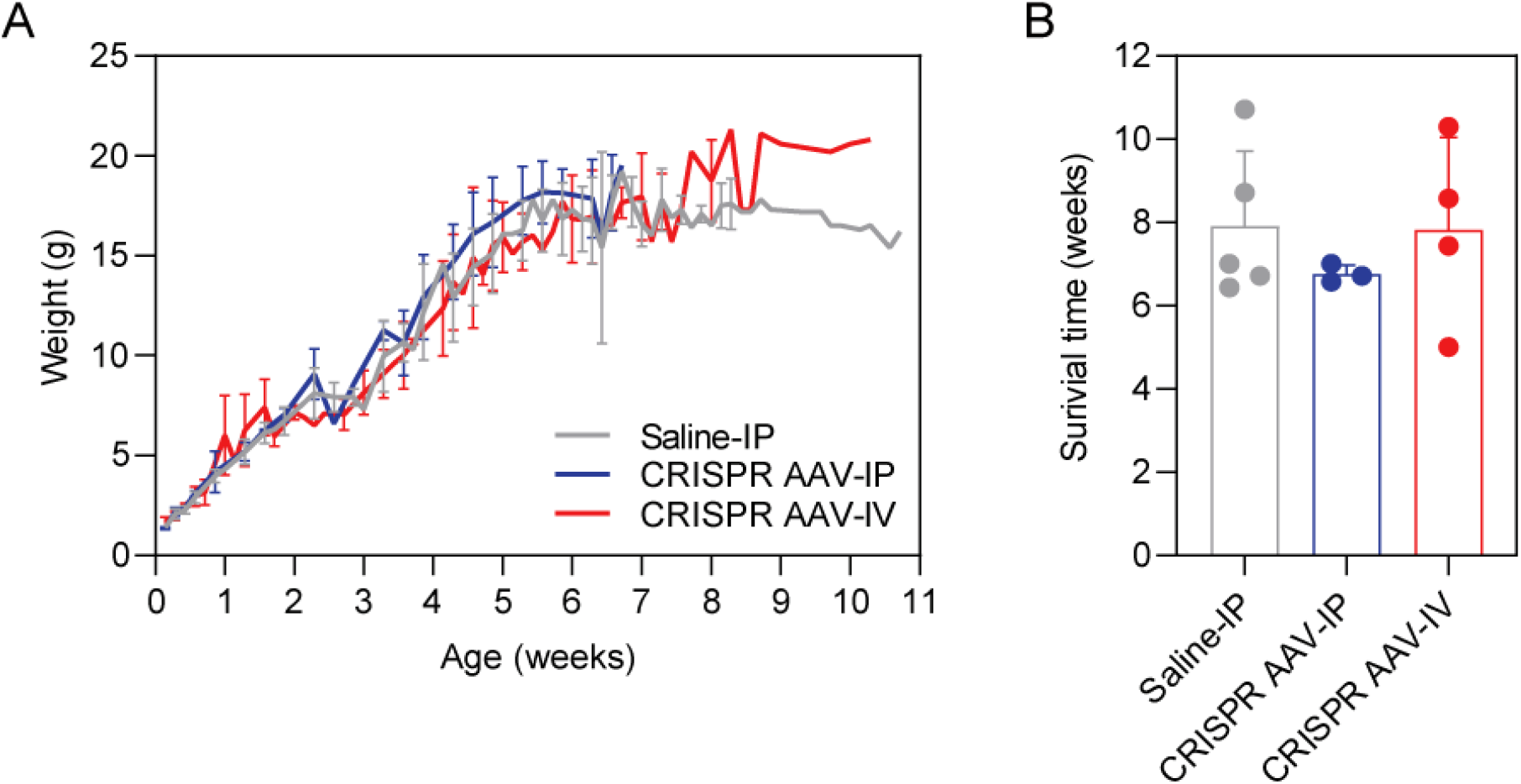
CRISPR AAV-treatment does not improve lifespan in dKO mice. dKO mice were treated with saline (*n*=5) or CRISPR AAV vectors by intraperitoneal (IP, *n*=3) or intravenous (IV, *n*=4) injection routes and weighed three times weekly until culling at the humane end point. (**A**) Animal body weight measurements over time, values are mean±SEM. (**B**) Age at humane endpoint (i.e. survival time), values are mean+SEM.

The biodistribution of AAV vectors delivered by either IP or IV delivery routes was assessed by measuring the number of vector genome copies (using primers against the SaCas9 transgene) per host genome by absolute quantification qPCR (**Figure S3**). AAV vectors were present at the highest levels in the heart and diaphragm tissues for both IP and IV injected animals, and consistent with gene editing outcomes. Notably, the levels of AAV vector genomes in the TA and triceps muscles was very low in the IP-treated mice, and between 26- and 95-fold lower than in the same tissues from IV-treated mice (**Figure S3**). These data suggest that the IV route of administration is important for enabling delivery to peripheral muscles.

### Non-uniform dystrophin localization in dystrophic muscle after CRISPR AAV treatment

We have previously reported on the importance of uniform dystrophin expression for correcting dystrophic pathology and preventing the release of circulating miRNA biomarkers using the *mdx*-*Xist*^Δhs^ mouse (that expresses dystrophin in a patchy manner as a consequence of skewed X-chromosome inactivation) ^52^. In the present study, some degree of within-fiber patchiness was observed in transverse sections (**Figures 3,5**, arrowheads), although this phenomenon is typically more easily seen in longitudinal sections. We therefore next sought to analyze the pattern of dystrophin expression in longitudinal sections of TA muscles derived from dKO mice that received CRISPR AAV treatment by either IP and IV route of injection (**Figure 7**). Such a patchy dystrophin distribution was observed for both treatment regimens, whereby adjacent regions of dystrophin-positive and dystrophin-negative sarcolemma were observed in the same myofiber (**Figure 7**).

**Figure 7.**
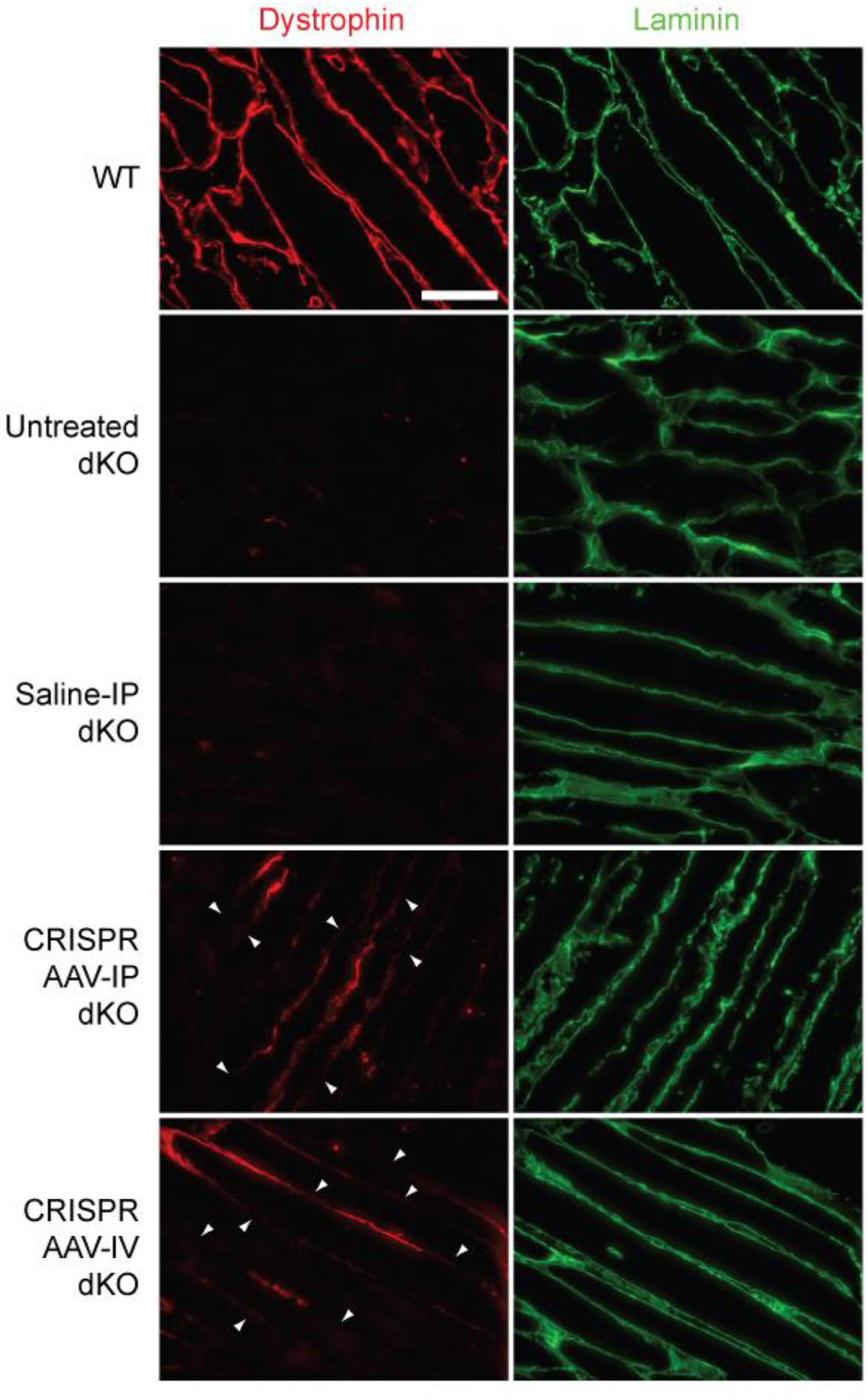
Dystrophin protein is re-expressed in a within-fiber patchy manner in dKO muscle after CRISPR AAV treatment. Representative dystrophin IF staining in longitudinal TA muscle sections of dKO mice treated with CRISPR AAV9 particles by IP or IV injection at P2. Untreated wild-type (WT), untreated dKO, and saline-treated dKO samples were analyzed in parallel as controls. (The pictured CRISPR-treated sections are derived from animals that express 1% and 3.3% of WT dystrophin for IP- and IV-treated animals respectively). Sections were co-strained for laminin to delineate myofiber boundaries. Images were taken at 20× magnification, and scale bars represent 100 μm. Arrowheads indicate regions of patchy sarcolemmal dystrophin staining.

### Detection of on-target, non-productive editing events by long-read sequencing

The patchy pattern of sarcolemmal dystrophin distribution is likely a consequence of the chimeric nature of the treated fibers, which contain both corrected (i.e. dystrophin-expressing) and uncorrected (i.e. non-dystrophin-expressing myonuclei). This scenario may arise as a result of incomplete gene editing, whereby the CRISPR/Cas9 machinery does not reach all myonuclei within a fiber. However, it is also expected that not all gene editing events will be repaired in a productive manner, meaning that the *Dmd* locus may be incorrectly edited in some myonuclei, such that they can never express functional dystrophin protein. To characterize the degree of non-productive gene editing, we performed targeted long-read sequencing of a ∼800 bp PCR amplicon covering *Dmd* exon 23 together with flanking intron sequence containing both sgRNA target sites using the PacBio sequencing platform. This analysis was performed in cardiac muscle, as gene editing was most effective in this tissue, and so these samples were expected to contain the most complex sequencing libraries. Sequencing reads were processed using a custom pipeline in order to assign them as unedited, productive edited (i.e. excision of *Dmd* exon23), and non-productive edited. Amplicons exhibiting non-productive editing were further classified as having (i) disruption of the 5ʹ sgRNA target site, (ii) disruption of the 3ʹ sgRNA target site, (iii) disruption of both sgRNA target sites, (iv) inversion of the excised fragment, and (v) AAV vector backbone integration (**Figure 8A**). The success of the classification was assessed by determining the amplicon size distributions for the classification bins (**Figure S4**).

**Figure 8.**
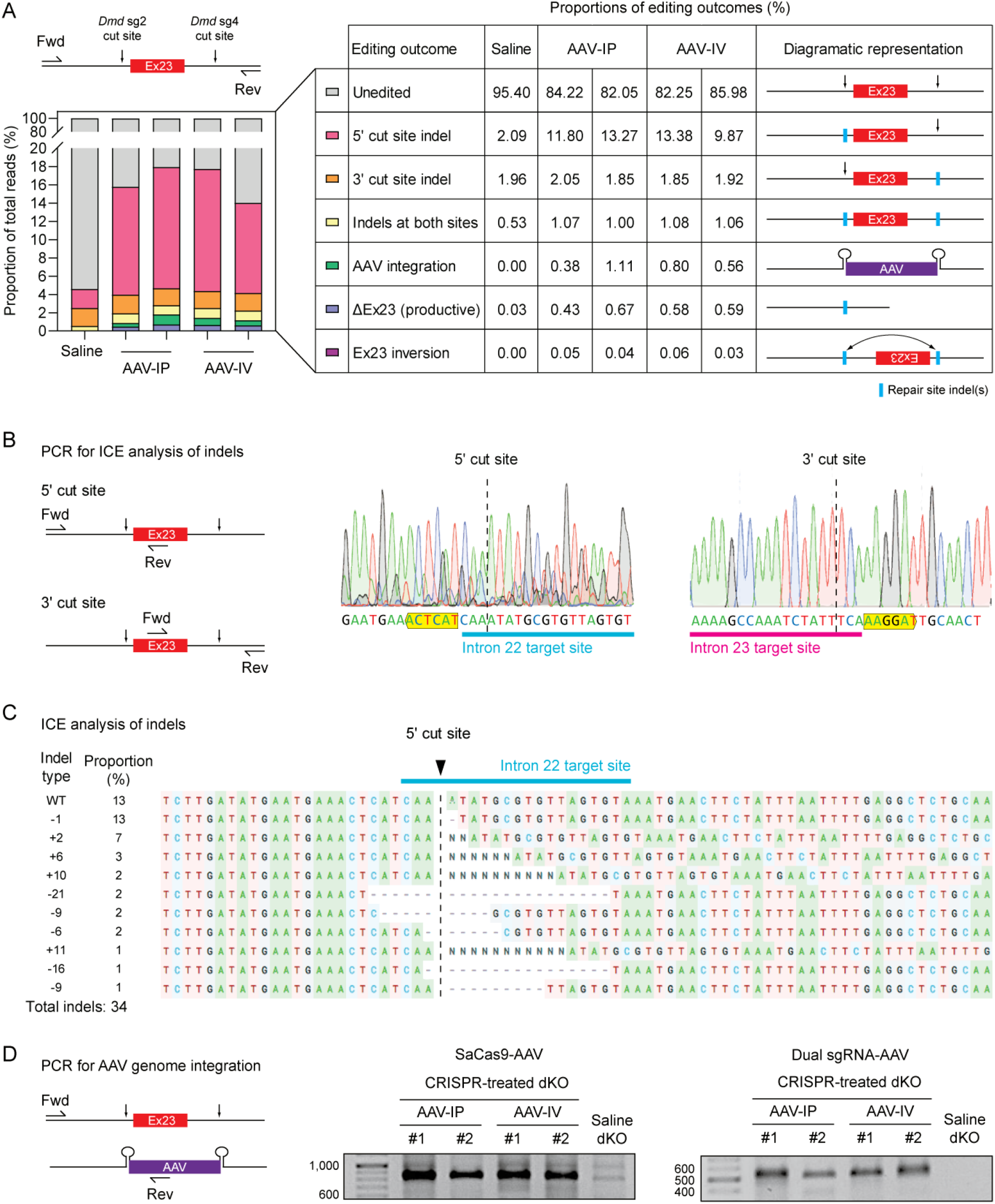
Detection of productive and non-productive editing outcomes by long-read DNA sequencing. DNA was harvested from dKO mice following treatment with CRISPR AAV (administered via either intraperitoneal or intravenous routes) and the edit region amplified by PCR (*n*=2, each). Saline-injected dKO tissue was collected in parallel as a control (*n*=1). Heart muscle was used in all cases as editing was found to be highest in this tissue. (**A**) PCR amplicons were analyzed by long-read next generation DNA sequencing and assigned to one of the following categories: unedited, indels at either sgRNA target site (or both), presence of integrated AAV backbone, productive excision of *Dmd* exon 23 (ΔEx23), and inversion of exon 23. The percentages of each editing outcome are indicated in the table. (**B**) The formation of indels was assessed at both 5ʹ and 3ʹ cut sites by PCR and Sanger sequencing. (**C**) The 5ʹ cut site was further analyzed using the Inference of CRISPR Edits (ICE) to characterize indel formation. (**D**) Integration of AAV backbones derived from both SaCas9-AAV and dual sgRNA-AAV vectors was confirmed in the treated animals by PCR at the 5ʹ cut site (**Figure S5**).

The majority (82-86%) of reads in the CRISPR-treated libraries were unedited, whereas productive excision of *Dmd* exon 23 was observed at a rate of 0.43-0.67% based on this analysis (**Figure 8A**). In contrast, only 0.03% of reads in the control sample were categorized as lacking *Dmd* exon 23. (Note: this analysis is expected to underestimate the degree of productive editing as the PCR reaction was optimized to preferentially amplify the full-length target).

A major limitation of the dual sgRNA targeting exon excision CRISPR strategy used in this study is that the success of this approach is predicated on simultaneous cleavage at both target sites. However, asymmetric cleavage, whereby one site is cut and repaired forming an indel before the neighboring site can be cut, will result in a non-productive gene editing outcome, and render the locus refractory to further editing. Asynchronous indel formation was observed at both the 5ʹ and 3ʹ sgRNA target sites, although the levels of such non-productive editing were much higher for the 5ʹ site (9.9-13.4%) (**Figure 8A**). While indels were detected at the 3ʹ sgRNA site, these were present at levels similar to the saline-treated control sample (i.e. background) (∼2%) (**Figure 8A**). These data suggest that CRISPR-mediated cleavage occurred much more efficiently at the 5ʹ sgRNA cut site than for the 3ʹ site. Similar results were obtained when these sites were analyzed by Sanger sequencing deconvolution using the ICE (Inference of CRISPR Edits) method (**Figure 8B,C**). We also detected reads in which indels were formed at both sgRNA sites, indicative of cleavage and repair, although the intervening *Dmd* exon 23 sequence was not excised. The relatively high percentage of asynchronous, non-productive indel formation at the 5ʹ sgRNA target site, is likely a major factor limiting the efficacy of this gene editing therapeutic approach.

A very small proportion of reads (∼0.03-0.06%) contained inversions of exon 23, whereby the region between the two cut sites was excised, and then re-inserted in the reverse orientation (**Figure 8A**).

The CRISPR-treated libraries also contained a substantial number (0.38-1.11%) of reads containing AAV-derived sequences (**Figure 8A**). These reads consisted of amplicons containing AAV vector backbone sequence from both vectors, and in both forward or reverse orientations. Notably, this rate of AAV integration events is likely to be an underestimate, as the PCR reaction is expected to be biased against the amplification of the larger template sequence resulting from full-length AAV backbone integration. Consistent with this notion, reads in the AAV integration classification bin exhibited shift towards a larger amplicon size which was not observed in other read categories, but was still much shorter than the full-length SaCas9-AAV genome (maximum detected size: 1,721 bp, **Figure S4**). The presence of AAV integration events was further confirmed by PCR of genomic DNA using a forward primer located in intron 22 and a reverse primer located in the CMV promoter of the AAV-SaCas9 vector, or in the U6 promoter of the AAV-dual sgRNA vector, as appropriate (**Figure 8D**). PCR amplicons of the expected size were only detected for CRISPR-treated animals, and Sanger sequencing of clones of these amplicons revealed the presence of AAV genome sequence as expected, although a variety of fragmentations were observed (**Figure S5**). Amplicon reads that were assigned to the *Dmd* exon 23 inversion or AAV integration event categories were extremely low in the saline-treated sample (i.e. 0% and 0.0022%, respectively). Taken together, these data are indicative of frequent, non-productive editing events in CRISPR-treated cardiac muscle.

## Discussion

Here we report the first study to demonstrate CRISPR-mediated rescue of dystrophin expression in the severely-affected dKO mouse model of DMD. Several other studies have employed a similar dual sgRNA CRISPR exon excision strategy for dystrophin restoration ^23–27, 31, 53^, although these have typically utilized the *mdx* mouse model which exhibits only a mild phenotype and almost normal lifespan ^23–25, 27^. While we observed dystrophin restoration in multiple dKO muscle tissues (**Figures 2**-**5, 7**), this was not sufficient to improve survival (**Figure 6**). In contrast, exon skipping using antisense oligonucleotides ^48, 54^ and the U7 snRNA system ^49^ has been shown to result in pronounced increases in lifespan from ∼8-10 weeks in untreated animals to between >6-months ^48, 54^ and ∼1 year, respectively ^49^ (although the levels of dystrophin expression reported in these reports were much higher than that observed in the present study).

We observed up to 4% of WT dystrophin protein levels in the heart and diaphragm muscles of dKO mice that received CRISPR AAVs by IP injection, and up to 2 and 3% in the heart and TA muscles of those treated by IV administration, respectively (**Figures 2,4**). Comparable levels of dystrophin have been previously reported in the hearts of CRISPR-treated *mdx* mice assessed at 3-weeks ^24^ and 8-weeks ^23, 25, 27^ post treatment (i.e. a similar time-frame to the limited lifespan of the dKO mouse). Notably, several *mdx* studies reported progressive increases in dystrophin protein levels at 6-months ^23^, 12-months ^25^, and 18-months ^31^ post treatment, with up to ∼20% restoration in the heart at the latest time point ^31^. It has been suggested that this accumulation of dystrophin is a consequence of its long half-life (>100 days) ^55, 56^, and of corrected myofibers experiencing a selection advantage over dystrophin-negative, unprotected fibers that remain sensitive to contractile damage and myonecrosis ^57^. In contrast, it is likely that the treated dKO animals used in the present study died before there had been sufficient time for corrected dystrophin protein to accumulate to levels above the threshold required for phenotypic correction. Indeed, the levels of dystrophin protein restoration that we observed fell short of the widely-accepted threshold of ∼15-30% that is believed to be required for phenotypic correction in *mdx* mice and DMD patients ^58, 59^, but are similar to the levels reported in human DMD patients treated with the FDA-approved exon skipping drug eteplirsen (i.e. treated patients expressed 0.93% dystrophin relative to healthy controls) ^18^.

The dose of each AAV9 vector that we utilized (1×10^11^ of SaCas9-AAV, and 5×10^11^ vg of dual sgRNA-AAV, a dose which is equivalent to ∼5×10^13^ and 2.5×10^14^ vg/kg respectively) is comparable to that used in published *mdx* studies at the time that this study was initiated ^23, 24, 27^. However, it has since emerged that both the dose of the AAV as well as the ratio of sgRNA:Cas9 are important determinants of gene editing success. The effect of increasing the dose of vectors carrying the CRISPR machinery was demonstrated by Bengtsson and colleagues, where the authors assessed co-excision of ‘hotspot’ exons 52 and 53 in the *mdx*^4c^*^v^* mouse (which harbors a nonsense mutation within exon 53) ^26^. The authors observed that widespread skeletal muscle restoration was only observed at the highest dose (i.e. 1×10^13^ vg of SpCas9-AAV and 4×10^12^ of sgRNA-AAV) ^26^. It is therefore possible that injection of higher doses of our vectors may result in improved dystrophin restoration and functional correction in the dKO mouse.

The dose of AAV required to achieve therapeutic dystrophin editing in patients is currently unknown, but based on pre-clinical studies is likely to be high. This raises the issue of whether sufficient quantities of virus can realistically be produced to treat all patients given current manufacturing demands/constraints. Furthermore, the feasibility of using high dose AAV regimens in patients has been called into question as important safety concerns have recently resurfaced. Specifically, in light of the deaths of child participants in a clinical trial for X-linked myotubular myopathy (NCT03199469) ^60^, and another child participant in a separate AAV trial for mucopolysaccharidosis type IIIA (NCT03612869) ^61^. Furthermore, during the preparation of this manuscript, Pfizer announced that a DMD patient treated with high dose AAV microdystrophin gene therapy has died, although few other details have been publicly disclosed at the time of writing ^62^. The dose of AAV administered could be reduced if more potent strategies are used, such as self-complementary AAV (scAAV). scAAV vectors form dsDNA hairpin structures, and thereby bypass the rate-limiting second strand synthesis step in the AAV life cycle, leading to accelerated transgene expression ^63, 64^. Zhang and colleagues observed a 70-fold improvement in editing efficiency in *mdx* mice with sgRNAs expressed from scAAV relative to conventional AAV at the same dose when applying a 1:10 ratio of AAV-Cas9 to AAV-sgRNA ^33^. However, we also performed additional injections using our vectors at this 1:10 ratio but observed negligible dystrophin restoration and no lifespan extension (data not shown). Alternative approaches to enhancing gene editing potency include improved capsids with stronger muscle tropism, de-targeting of vectors from the liver, higher Cas9/sgRNA expression, and better sgRNA design.

Analysis of longitudinal sections of CRISPR-treated dKO mouse skeletal muscle tissue revealed a ‘within-fiber’ patchy pattern of dystrophin restoration following CRISPR treatment (**Figure 7**), a phenomenon that we had previously predicted ^34, 52, 65^. The observed patchiness is consistent with the myonuclear domain hypothesis, whereby each myonucleus serves its proximal cytoplasmic territory ^66^. To our knowledge, no other DMD CRISPR study has analyzed dystrophin immunostaining in longitudinal muscle sections. The functionality of muscle relies on coordinated contraction and force dissipation along the length of the myofiber, and as such, incomplete sarcolemmal dystrophin coverage following gene editing could result in a failure to achieve functional correction. The role of dystrophin is to protect myofibers from contractile damage ^67^, so in the case of patchy patterns of sarcolemmal dystrophin expression, molecular pathological events could still initiate at unprotected dystrophin-negative regions within the myofiber. These observations may partially explain why no lifespan extension was observed in this study. Interestingly, Torelli *et al*. recently reported incomplete sarcolemmal dystrophin coverage in human patients with severe BMD and intermediate muscular dystrophy (i.e. disease severity greater than typical BMD, but less severe than typical DMD) ^68^.

A similar patchy dystrophin pattern was observed in the *mdx*-*Xist*^Δhs^ mouse which expresses varying levels of dystrophin as a result of skewed X-chromosome inactivation ^52, 69^. The myofibers of both the *mdx*-*Xist*^Δhs^ mouse and CRISPR-treated dKO mice are heterokaryons that contain both dystrophin-expressing, and non-dystrophin-expressing myonuclei, which likely explains the similarity between these model systems. These findings strongly suggest that dystrophin mRNA and protein are subject to a degree of spatial restriction, and are not free to diffuse throughout the sarcoplasm and sarcolemma, respectively. While the levels of dystrophin restoration in our CRISPR-treated animals was low, patchiness in the *mdx-Xist*^Δhs^ model was observed even with >40% of WT dystrophin expression levels ^52^. We therefore propose that the *mdx-Xist*^Δhs^ mouse is a putative model that simulates the outcome of CRISPR-mediated dystrophin correction over a range of effectiveness levels. In contrast, we have observed that antisense oligonucleotide-mediated exon skipping with peptide-PMO conjugates results in within-fiber uniform dystrophin distribution that is independent of dose (**see related manuscript**).

Notably, van Putten and colleagues generated a variant dKO model crossed with the *Xist*^Δhs^ mouse (i.e. *mdx*/utrn^-/-^/*Xist*^Δhs^) ^70^. In this model, an increase in survival was observed in mice expressing as little as ∼4% WT dystrophin levels ^70^. Crucially, these mice experienced the restorative benefits of bodywide expression of full-length dystrophin protein from birth, whereas CRISPR treatment in the present study resulted in the generation of a BMD-like dystrophin, with a partial internal truncation and attenuated functionality ^71^, that was differentially restored across muscle tissues. Additionally, the *mdx*/utrn^-/-^/*Xist*^Δhs^ mice are necessarily female (as variable dystrophin expression is dependent on skewed X-chromosome inactivation), which is a further potentially confounding factor.

Instances in which Cas9-mediated cleavage occurs and the DSB is repaired in a non-productive manner (i.e. corruption of one or more of the sgRNA target sites ^53^, inversion of excised region ^72^, or integration of sequence from the AAV vector genome ^25, 32, 73^) are likely to contribute to myofiber heterogeneity as the resulting myonuclei are incapable of producing dystrophin, or being re-cut by Cas9. Such non-productive editing events were readily detected in the heart tissue of CRISPR-treated dKO animals (**Figure 8**). In particular, we observed a relatively high proportion of indel formation at the 5ʹ sgRNA cut site, indicative of asynchronous cleavage and repair. Such nuclei are now rendered refractory to further editing as the introduction of an indel disrupts the sequence of the sgRNA binding site. This observation of differential sgRNA cutting efficiency was somewhat surprising, given that initial *in vitro* screening suggested that both guides were effective (**Figure 1**). These data are also suggestive of differential cleavage potential for the 5ʹ and 3ʹ sgRNAs, which is likely limiting the overall dystrophin restoration efficacy of this therapeutic approach.

Importantly, while there is a possibility of CRISPR-induced lesions being resolved in a non-productive manner, the generation of myofiber heterokaryons is inevitable. In other words, even if DNA cleavage is achieved in every single myonucleus, some of these will be non-productively repaired, meaning that the treated fibers must necessarily consist of both dystrophin-expressing and non-dystrophin-expressing nuclei. An alternative to entire exon excision is the use of a single-cut CRISPR strategy to disrupt splice acceptor sites and putative exonic splicing enhancer motifs in the gDNA. As a result, the spliceosome is no longer able to recognize these splicing signals, thereby achieving permanent ‘exon skipping’. This strategy has been utilized in *mdx* mice ^26, 29, 32, 74^, as well as a canine model of dystrophy ^30^, with high levels of gene editing success. For this approach, the formation of an indel at the cut site is expected to result in restoration of dystrophin expression, whereas seamless repair without an indel regenerates the sgRNA target site. As such, the single-cut CRISPR strategy may be superior to the dual cut approach owing to the simplicity of design and propensity for a greater number of productive editing outcomes. However, AAV genome integration is still possible at the DSB site for the single-cut approach, and the single-cut strategy cannot be leveraged for multi-exon excision.

In conclusion, CRISPR-mediated dystrophin restoration in the severely-affected dKO mouse did not reach a sufficient level to achieve an improvement in survival. Future developments are required to increase the overall editing efficiency of the CRISPR AAV dual-cut strategy employed in this study including optimization of delivery to a broad range of muscle tissues, particularly the heart and diaphragm, and reducing the likelihood of non-productive editing outcomes. This study highlights the importance of uniform dystrophin expression in the successful implementation of genetic therapies for DMD.

## Methods

### Plasmids and AAV Production

For cell culture studies, guide RNAs were expressed using pTZ-U6sgRNA(SaCas9) vectors that were modified to encode specific *Dmd*-targeting sequences (**Table S1**) by Golden Gate Assembly ^75^. SaCas9 was expressed using pX601-AAV-CMV::NLS-SaCas9-NLS-3xHA-bGHpA;U6::BsaI-sgRNA) which was a gift from Feng Zhang (Addgene plasmid #61591; http://n2t.net/addgene:61591; RRID:Addgene_61591) ^41^.

AAV2/9 vectors were generated by Atlantic Gene Therapies (University of Nantes, Institut de Recherche Thérapeutique, Nantes, France) using pX600 (SaCas9-AAV) (Addgene, Watertown, MA, USA) and pAAVio-2x.sgRNA (*Dmd* dual sgRNA-AAV).

### Cell Culture

10T1/2 mouse fibroblasts cells (a kind gift from Peter K. Vogt) were cultured in DMEM (supplemented with 10% FBS and 1× antibiotic/antimycotic (all Thermo Fisher Scientific, Abingdon, UK) and maintained in a humidified incubator at 37°C with 5% CO_2_. Cells were seeded at 1×10^5^ cells per well in a 24 well multiplate and transfected with 800 ng plasmid DNA per well (400 ng of SaCas9 expression plasmid and 200 ng of each sgRNA plasmid) using Lipofectamine 2000 (Thermo Fisher Scientific) as according to manufacturer’s instructions, and harvested 48 hours later.

### Animal Studies

All procedures were authorized by the UK Home Office (project license 30/2907) in accordance with the Animals (Scientific Procedures) Act 1986. Animals were housed under 12-hour light/12-hour dark conditions, with access to food and water *ad libitum*.

Utrophin/dystrophin double knockout (dKO) mice were obtained in the F1 generation by breeding with *Utrn^tm1Ked^Dmd^mdx^*/J mice (JAX stock #019014; Jackson Laboratory, Bar Harbour, ME, USA) which are heterozygous for the utrophin knockout mutation and homozygous (females)/hemizygous (males) for the *mdx* dystrophin knockout mutation ^46^. Wild-type C57BL/10 (C57BL/10ScSn) and dystrophic *mdx* (C57BL/10ScSn-*Dmd^mdx^*/J) mice were also used as controls as appropriate.

dKO mice were treated at P2, using either the intraperitoneal (IP, *n* = 3) or intravenous (IV via the facial vein, *n* = 4) injection routes, with a 20 µl of a mixture of the two AAV vectors in 0.9% sterile saline. The vectors were formulated as a 1:5 SaCas9:sgRNA mixture consisting of 1×10^11^ vg SaCas9-AAV and 5×10^11^ vg *Dmd* dual sgRNA-AAV. Control dKO mice were treated with 0.9% sterile saline (*n*=5).

Mice were weighed three times a week, and sacrificed at the humane end point (which was determined by rapid labored breathing, and reduced mobility leading to difficulty accessing food and water) by increasing CO_2_ concentration. Serum was collected from the jugular vein immediately after termination at the humane endpoint, using Microvette CB 300 Blood Collection tubes (Sarstedt, Nümbrecht, Germany) as described previously ^76^. Serum was allowed to clot and separated by centrifugation at 10,000 *g* for 5 min at 4 °C. The heart, diaphragm, TA, and triceps muscles were harvested by macrodissection. Tissues were mounted onto cork discs using O.C.T. Compound Mounting Media for Cryotomy (VWR, Lutterworth, UK) and snap frozen in dry ice cooled isopentane. Tissue samples were stored at -80°C prior to downstream processing. Tissues from *mdx* and WT mice were collected in parallel for the purposes of generating western blot standard curves and control sections, as appropriate.

### Genomic DNA Analysis

Genomic DNA was extracted from cells using KAPA Express Extract (Kapa Biosystems, Feltham, UK) and from tissues using the DNeasy Blood & Tissue Kit (Qiagen, Manchester, UK), according to the manufacturer’s instructions. PCR primers were designed to amplify a sequence that spans the targeted deletion site (**Table S2**). For cell culture samples, PCR was performed using KAPA 2G Fast HotStart ReadyMix (Kapa Biosystems) with an initial denaturation phase at 95°C for 3 minutes followed by 35 cycles of denaturing (95°C for 15 seconds), annealing (53°C for 15 seconds) and extension (72°C for 30 seconds). For *in vivo* samples, PCR was performed using KAPA2G Robust HotStart ReadyMix (Kapa Biosystems) with PCR conditions as above (with the annealing temperature modified to 55°C). PCR products were separated by agarose gel electrophoresis. PCR amplicons were purified and gene editing confirmed by Sanger sequencing (Source BioScience, Nottingham, UK). Individual amplicons were cloned using the TOPO TA cloning for sequencing kit (Thermo Fisher Scientific). Sanger sequencing chromatogram deconvolution was performed using the ICE online tool ^77^.

### Quantitative PCR

Reverse transcriptase-quantitative polymerase chain reaction (RT-qPCR) was conducted following the MIQE (minimum information for publication of quantitative real-time PCR experiments) guidelines ^78^. Total RNA was isolated from muscle tissues (∼100 µm sections, or 1/6^th^ of the diaphragm) using a Maxwell RSC Instrument and Maxwell RSC simplyRNA Tissue Kit (both Promega, Southampton, UK) and complementary DNA (cDNA) was reverse-transcribed using the High-Capacity cDNA Reverse Transcription Kit (Thermo Fisher Scientific) according to the manufacturer’s instructions. Quantitative PCR (qPCR) was carried out using TaqMan probe-based assays to determine the level of *Dmd* exon 23 ‘skipping’ as a consequence of transcription from the edited *Dmd* locus. qPCR reactions were performed using TaqMan Universal PCR Master Mix (Thermo Fisher Scientific) and the StepOnePlus Real-Time PCR System (Applied Biosystems) with the following cycling conditions: 95 °C for 10 minutes, followed by 40 cycles of 95 °C for 15 seconds and 60 °C for 1 minute.

Absolute quantification was performed by comparing samples to standard curves comprised of serial dilutions of the full-length and ΔExon23 *Dmd* DNA target templates (IDT, Leuven, Belgium). The degree of exon 23 excision was determined by calculating the percentage of ΔExon23 *Dmd* transcripts relative to the total (i.e. full-length + ΔExon23 *Dmd* transcripts). All primer sequences are listed in **Table S2**.

AAV vector genome copies were determined by absolute quantification qPCR using primers designed to amplify the SaCas9 transgene (**Table S2**). Samples were compared to serial dilutions of plasmid DNA in order to calculate copy numbers. To measure host genome copy numbers, a commercial TaqMan assay designed to amplify the *Actb* gene (Mm00607939_s1, Applied Biosystems) was used, and amplification results compared against a serial dilution of mouse genomic DNA of known mass. qPCR was performed as described above, except for SaCas9 mRNA quantification, where Power SYBR Green Master Mix (Thermo Fisher Scientific) was used.

### Serum miRNA Analysis

Extracellular miRNAs were analyzed by small RNA TaqMan RT-qPCR as described previously ^76, 79^. Briefly, RNA was extracted from 50 µl of serum using TRIzol LS (Thermo Fisher Scientific) according to manufacturer’s instructions, with minor modifications. RNase-free glycogen (Roche) was used as an inert carrier to maximize RNA recovery, and a synthetic oligonucleotide spike-in control (cel-miR-39, 5ʹ-UCACCGGGUGUAAAUCAGCUUG-3ʹ, IDT) was added at the phenolic extraction stage, to enable later data normalization. RNA was re-suspended in 30 µl of nuclease-free water (Thermo Fisher Scientific) and 10 µl of each sample used for reverse transcription using the TaqMan MicroRNA Reverse Transcription Kit (Thermo Fisher Scientific) according to manufacturer’s instructions. Thermal cycling was performed as described above. Data were normalized to cel-miR-39 and data analyzed using the Pfaffl method ^80^. miRNA TaqMan assays are described in **Table S3**.

### Western Blot

Total protein was isolated from muscle tissues (∼100 8 μm sections, or 1/6^th^ of the diaphragm) using modified RIPA lysis buffer (10% SDS, 50 mM tris (pH 8), 150 mM NaCl, 1% NP-40, 0.5% sodium deoxycholate), supplemented with 1× cOmplete protease inhibitor cocktail (Merck, Feltham, UK). Samples were homogenized using a Precellys 24 Tissue Homogenizer (Bertin Technologies, Paris, France) (4 cycles at 5,000 rpm for 30 seconds). 20 μg of total protein lysate for each sample was separated by SDS-PAGE, using NuPAGE 3-8% Tris-Acetate 1 mm 10-well gels (Thermo Fisher Scientific) run at 150 V for 90 minutes in 1× NuPAGE Tris-Acetate SDS Running Buffer (Thermo Fisher Scientific). Proteins were electrotransferred onto an Immobilon-fl polyvinylidene difluoride (PVDF) membrane (Merck) at 30 V for 1 hour followed by100 V for a further hour at 4°C in 1× NuPAGE Transfer Buffer (Thermo Fisher Scientific) supplemented with 0.1 g/l of SDS (Sigma-Aldrich, MO, USA) and 10% methanol. Membranes were blocked in Intercept (PBS) Blocking Buffer (Sigma-Aldrich) for 30 minutes at room temperature, and then incubated with primary antibodies in Intercept Blocking Buffer supplemented with 0.1% Tween-20 (Sigma-Aldrich) at 4°C overnight. Membranes were subsequently washed with PBS supplemented with 0.1% Tween-20 (PBST), and incubated with secondary antibodies (Thermo Fisher Scientific) in Blocking Buffer for 1 hour at room temperature. After further washing in PBST, blots were developed using Clarity Western ECL Substrate (Bio-Rad, Watford, UK) and imaged using a LI-COR Biosciences Odyssey Fc instrument (LI-COR, NE, USA). Protein standards were generated by combining dystrophin-positive (WT) and dystrophin-negative (*mdx*) protein lysates in a series of ratios, and were run in parallel to facilitate dystrophin quantification in treated samples. Vinculin was used as a negative control. Details of all antibodies are shown in **Table S4**.

### Immunofluorescence

Eight-micrometer transverse or longitudinal sections of heart, diaphragm, TA, and triceps tissues were cryosectioned, transferred to Superfrost Microscope Slides (Thermo Fisher Scientific), and air dried prior to storage at -80°C. Sections were soaked in PBS for 10 minutes before incubation in blocking solution (20% fetal calf serum (FCS, Thermo Fisher Scientific), and 20% normal goat serum (NGS, MP Biomedicals, Eschwege, Germany), in PBS) for an hour at room temperature. Sections were then incubated with primary antibodies in blocking solution for 2 hours at room temperature. Sections were washed three times with PBS and then incubated with fluorescent secondary antibodies in PBS for 1 hour at room temperature. Slides were again washed three times with PBS and mounted with Vectashield Hard Set mounting medium with DAPI H-500 (2BScientific, Oxfordshire, UK). Sections were imaged using a Leica DMIRB Inverted Modulation Contrast Microscope using the MetaMorph imaging software (Leica Biosystems, Newcastle upon Tyne, UK). Images were processed using ImageJ software. Details of all antibodies are shown in **Table S4**.

### Long-Read Amplicon Sequencing

Genomic DNA was harvested from cardiac tissue of dKO mice treated with CRISPR AAVs either by IP or IV injection (*n*=2, each), as well as saline-treated control mice (*n*=1), using the DNeasy Blood and Tissue kit (Qiagen, Manchester, UK). The region spanning the CRISPR target site was amplified by PCR. Briefly, PCR was carried out on 100 ng of gDNA per sample using Phusion High-Fidelity DNA Polymerase (New England Biolabs, Hitchin, UK) with an initial denaturation phase (98°C for 30 seconds), and then 35 cycles of denaturing (98°C for 10 seconds), annealing (53°C for 15 seconds) and extension (72°C for 30 seconds). PCR products were purified using the GeneJET PCR purification kit (Thermo Fisher Scientific). Sample concentrations were determined using the QuantiFluor ONE dsDNA System and a Quantus Fluorometer (both Promega, Southampton, UK). Equimolar amounts of each sample were pooled, and long-read next-generation sequencing using the PacBio platform carried out as a service by Genewiz (South Plainfield, NJ, USA). PCR primers with internal 8-mer barcodes used for de-multiplexing (bold sequence) are listed in **Table S2.**

The barcoded circular consensus reads were demultiplexed, and asymmetric barcodes were removed using the LIMA software package v2.4.0 (https://github.com/pacificbiosciences/barcoding/). The resulting demultiplexed FASTQ files were inspected with FastQC v.0.11.9 (http://www.bioinformatics.babraham.ac.uk/projects/fastqc/), then quality filtered with BBDuk –part of the BBTools software toolkit v.38.93 (https://sourceforge.net/projects/bbmap/), with the maq parameter set to Q20. Reads shorter than 400 bases were discarded, resulting in more than 20,000 high quality reads per sample for all subsequent analyses. Sequence classification based on presence or absence of sequence features was performed with the BBDuk and BBDuk2 packages, which are designed to filter based on look up of *k*-mers. *k*-mer length was set to 25 bases, with a minimal edit distance, allowing for accurate filtering of reads which may contain minor sequencing errors. For example, a read was classified as AAV-containing, if it contained any 25 nucleotide contiguous sequence from the AAV reference, with up to 2 mismatches or indels allowed. The *k*-mer and edit distance parameters were optimized against the untreated sample, in order to minimize incorrect classification. Furthermore, at each sequence classification step, the resulting FASTQ files were aligned to the mouse chromosome X reference sequence (mm10), using the minimap2 SMRT wrapper for Pacbio data: pbmm2 v1.3.0 ^81^. The aligned reads were inspected using IGV v2.11.2, which supports 3^rd^ generation sequencing. FASTQ file intersections based on common read identifiers were generated using seqkit v2.1.0 ^82^. Raw sequencing data are deposited in NCBI SRA: PRJNA789451.

### Statistics

Statistical analysis was carried out using GraphPad Prism 9.0 software.

## Acknowledgements

B.H. and K.C. are supported by studentships from the Clarendon Fund. N.S. is supported by a studentship from Wave Life Sciences. Work in the laboratory of MJAW is supported by grants from the UK Medical Research Council.

