## Supplementary Information for "Non-uniform dystrophin re-expression after CRISPR-mediated exon excision in the dystrophin/utrophin double-knockout mouse model of DMD"

| ID | Target Sequence (5' to 3') |
|---|---|
| **Dmd-ex23-sgRNA-1** | TTATTTACCTTCTTCTTGAT |
| **Dmd-ex23-sgRNA-2** | CAAATATGCGTGTTAGTGTA |
| **Dmd-ex23-sgRNA-3** | AGTCCTTCAAAGATATTGAT |
| **Dmd-ex23-sgRNA-4** | CAAAAGCCAAATCTATTTCA |

**Table S1**

**Targets for sgRNA sequences.**

| ID | Sequence (5' to 3') |
|---|---|
| **gDNA Analysis** | |
| **Dmd-check-Fwd** | GCCTAAATGTCTTAATAATGTTTCAC |
| **Dmd-check-Rev** | GCTGTGAGCTAAATCATATCTACA |
| **Exon Skipping RT-qPCR** | |
| **qExon22-24-Fwd** | CTGAATATGAAATAATGGAGGAGAGACTCG |
| **qExon22-24-Rev** | CTTCAGCCATCCATTTCTGTAAGGT |
| **qExon22-24-Probe** | /5FAM/ATGTGATTC/ZEN/TGTAATTTCC/3IABkFQ/ |
| **qExon23-24-Fwd** | CAGGCCATTCCTCTTTCAGG |
| **qExon23-24-Rev** | GAAACTTTCCTCCCAGTTGGT |
| **qExon23-24-Probe** | /5HEX/TCAACTTCA/ZEN/GCCATCCATTTCTGTAAGGT/3IABkFQ/ |
| **DNA qPCR** | |
| **qSaCas9-Fwd** | CCAACGCCGATTTCATCTTC |
| **qSaCas9-Rev** | GATCTCTTTGTACTCCTGCTCG |
| **Long-read amplicon sequencing** | |
| **A701_Dmd_Check_Fwd** | **ATCACGAC**GCCTAAATGTCTTAATAATGTTTCAC |
| **A501_Dmd_Check_Rev** | **AAGGTTCA**GCTGTGAGCTAAATCATATCTACA |
| **A702_Dmd_Check_Fwd** | **ACAGTGGT**GCCTAAATGTCTTAATAATGTTTCAC |
| **A502_Dmd_Check_Rev** | **ACTTAGCA**GCTGTGAGCTAAATCATATCTACA |
| **A703_Dmd_Check_Fwd** | **CAGATCCA**GCCTAAATGTCTTAATAATGTTTCAC |
| **A503_Dmd_Check_Rev** | **AGAGAACA**GCTGTGAGCTAAATCATATCTACA |
| **A704_Dmd_Check_Fwd** | **ACAAACGG**GCCTAAATGTCTTAATAATGTTTCAC |
| **A504_Dmd_Check_Rev** | **GTGTCTTA**GCTGTGAGCTAAATCATATCTACA |
| **A705_Dmd_Check_Fwd** | **ACCCAGCA**GCCTAAATGTCTTAATAATGTTTCAC |
| **A505_Dmd_Check_Rev** | **TCGATTAG**GCTGTGAGCTAAATCATATCTACA |

**Table S2**

**List of primer sequences used in this study.**

Exon skipping RT-qPCR assay probes include 5′ terminal fluorophores (either FAM or HEX), 3′-terminal Iowa Black fluorescence quencher, and contain internal ZEN modifications. Barcode regions in the primers used for long-read amplicon sequencing are highlighted in bold.

| Target | Product ID | Detection Channel |
|---|---|---|
| **Small RNA TaqMan RT-qPCR** | | |
| **mmu-miR-1a-3p** | 002222 | FAM |
| **mmu-miR-133a-3p** | 002246 | FAM |
| **mmu-miR-206-3p** | 000510 | FAM |
| **mmu-miR-483-3p** | 002560 | FAM |
| **cel-miR-39** | 000200 | FAM |
| **DNA qPCR** | | |
| ***Actb*** | Mm00607939_s1* | VIC |

**Table S3**

**List of TaqMan assays used in this study.**

All products were obtained from Thermo Fisher Scientific.

| Target protein | Host | Product ID | Manufacturer | Dilution |
|---|---|---|---|---|
| **Western Blot** | | | | |
| **Dystrophin (DMD)** | Mouse mAb | NCL-DYS1 | Leica Biosystems | 1:100 |
| **Vinculin (VCL)** | Mouse mAb | V9131 | Sigma-Aldrich/Merck | 1:200 |
| **Anti-mouse IgG, HRP-linked** | Horse | 7076 | Cell Signaling Technology | 1:5,000 |
| **Immunofluorescence** | | | | |
| **Dystrophin (DMD)** | Rabbit pAb | ab15277 | Abcam | 1:1,000 |
| **Laminin subunit alpha 2 (LAMA2)** | Rat mAb | L0663 (clone: 4H8-2) | Sigma-Aldrich/Merck | 1:1,000 |
| **Anti-rabbit IgG Alexa Fluor 594** | Goat | ab150080 | Abcam | 1:500 |
| **Anti-rat IgG Alexa Fluor 488** | Goat | ab150157 | Abcam | 1:500 |

**Table S4**

**Antibodies used in this study.**

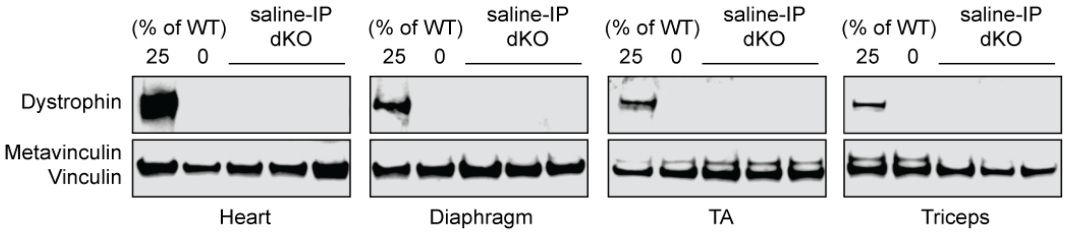


**Figure S1**

**Dystrophin protein is undetectable by western blot in saline-treated dKO mouse tissues.**

Western blot for dystrophin in 20 μg of total protein obtained from the heart, diaphragm, TA, and triceps muscles. Positive control samples contain a mixture of 25% WT (C57/BL10) with 75% dystrophin-deficient *mdx* protein. Negative control samples contain *mdx* protein only. The remaining samples consist of dKO animals treated with saline by IP injection. Vinculin was included as loading control.

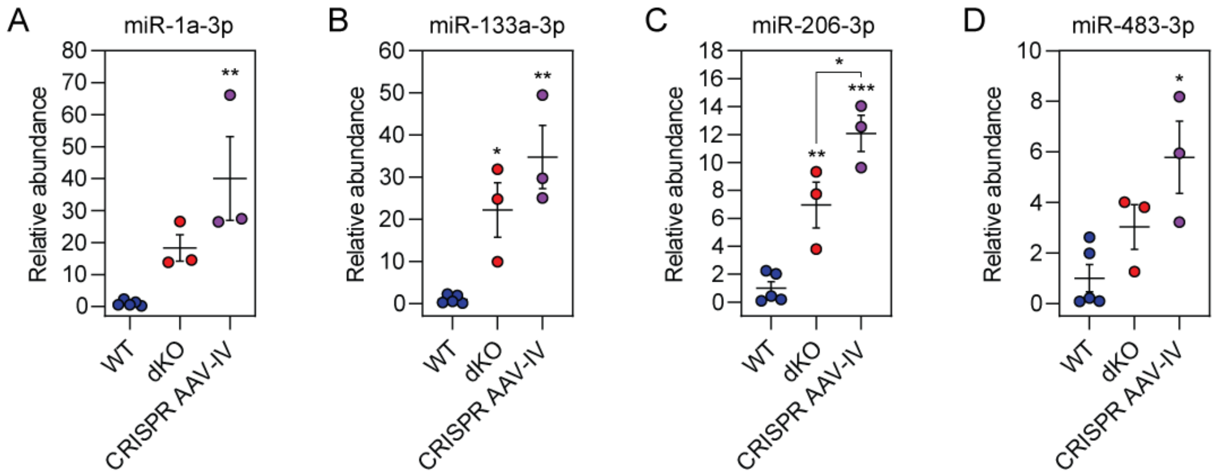


**Figure S2**

**miRNA biomarkers are not restored to wild-type levels in CRISPR AAV-treated dKO mouse serum.**

dKO mice were treated by intravenous (IV) injection via the facial vein at postnatal day two (P2) with $1\times10^{11}$ vg of SaCas9-AAV and $5\times10^{11}$ vg of dual sgRNA-AAV and sacrificed at the humane end point. Serum was harvested from the animals immediately postmortem (*n*=3) and RNA extracted. Serum RNA was analyzed for (**A**) miR-1a-3p, (**B**) miR-133a-3p, (**C**) miR-206-3p, and (**D**) miR-483-3p by small RNA TaqMan RT-qPCR. Serum from un-injected wild-type (WT) (*n*=5) and dKO (*n*=3) animals served as controls (10-weeks-old). Values are mean±SEM. Statistically significant differences were tested by one-way ANOVA with Bonferroni *post hoc* test, *$P<0.05$, **$P<0.01$, ***$P<0.001$.

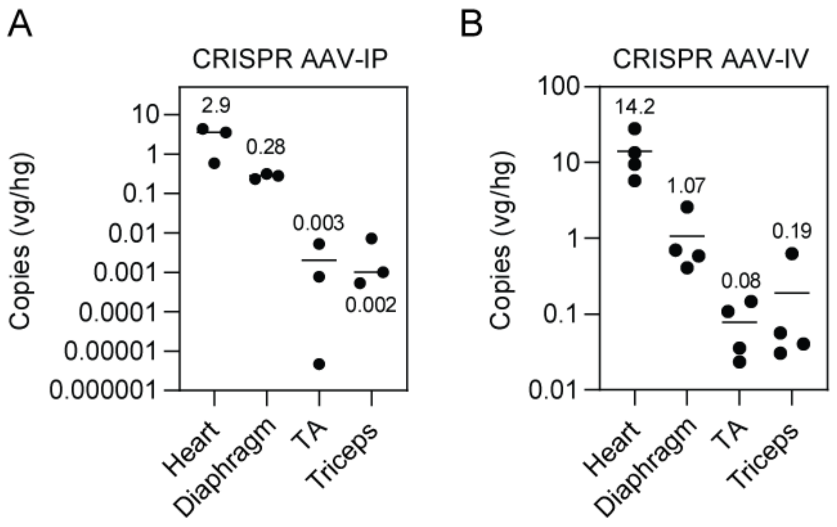


**Figure S3**

**Biodistribution of SaCas9 vector genomes.**

DNA was extracted from CRISPR AAV-treated dKO mice for (**A**) intraperitoneal (IP, *n*=3), and (**B**) intravenous (IV, *n*=4) routes of administration. Absolute quantification qPCR was used to detect SaCas9-AAV vector genome (vg) copies using primers against the SaCas9 transgene and data normalized to the copy number of host genomes (hg) measured using qPCR primers for *Actb*. Mean copy numbers are indicated by horizontal bars, and the mean value shown next to each column.

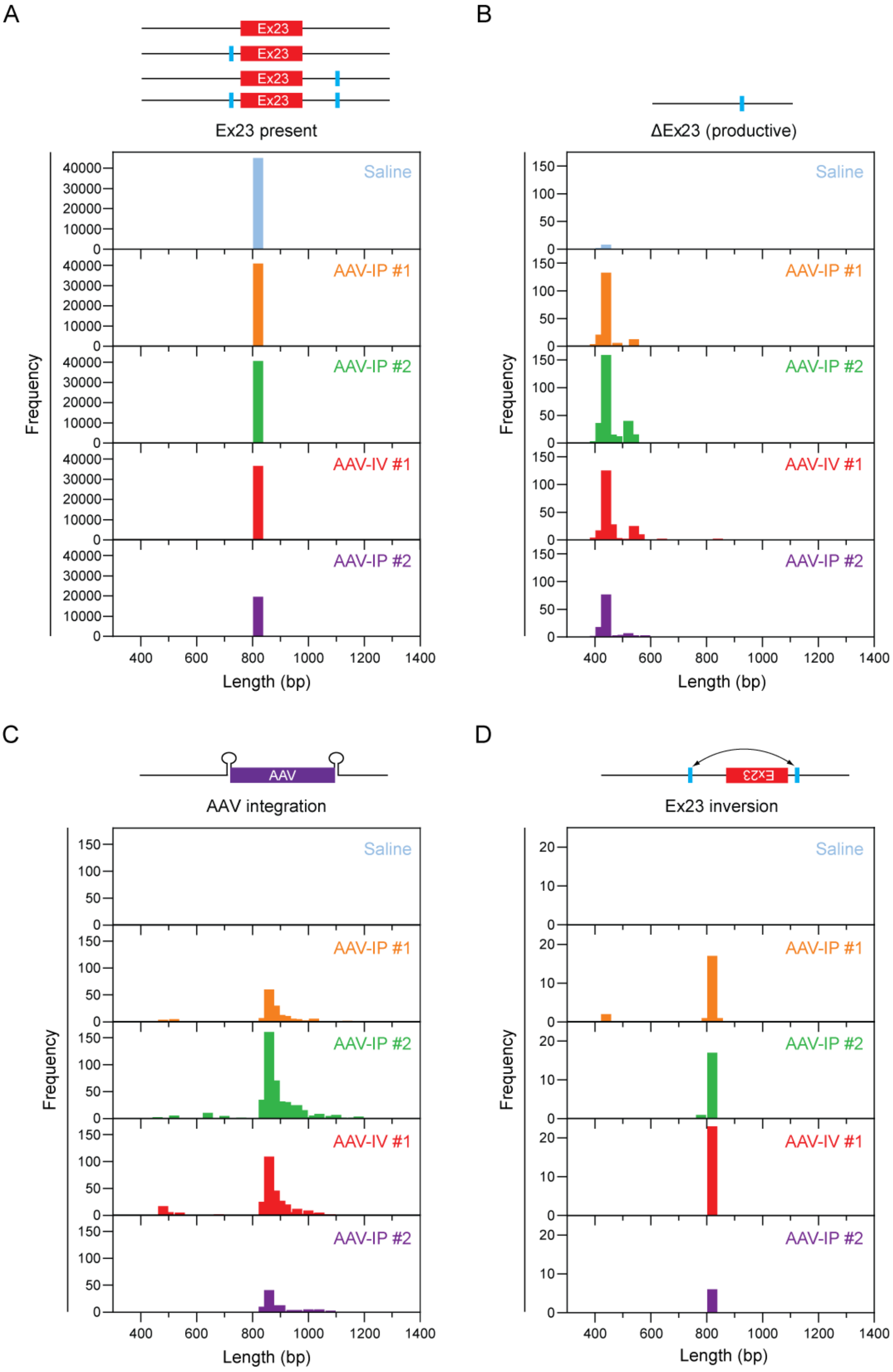
A
Ex23
Ex23 present
B
ΔEx23 (productive)
C
AAV
AAV integration
D
Ex23
Ex23 inversion
Saline
AAV-IP #1
AAV-IP #2
AAV-IV #1
AAV-IP #2
Frequency
Length (bp)

**Figure S4**

**Size distributions for read classification groups following long-read amplicon sequencing.** Sequenced amplicons from CRISPR-treated dKO heart samples, or saline control, were assigned to separate categories (**Figure 8**) and read length size distributions determined for (**A**) unedited sequence/indels present at the sgRNA cut site(s), (**B**) ΔEx23 (productive editing), (**C**) presence of integrated AAV-derived sequences, and (**D**) *Dmd* Ex23 present in the reverse orientation (i.e. Ex23 inversion). Schematic representations of possible amplicons are shown for each category.

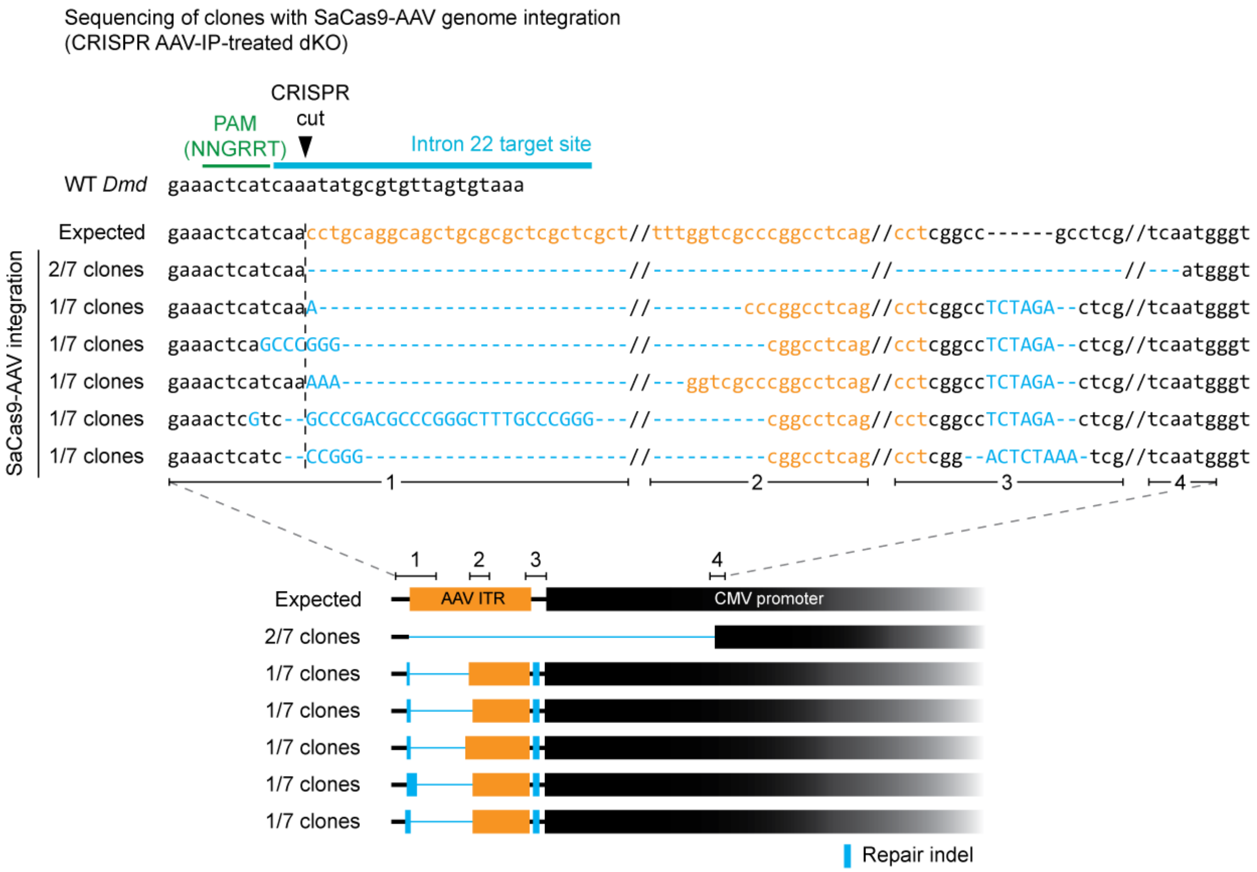


**Figure S5**

**Detection of SaCas9-AAV vector backbone at the 5′ CRISPR cut site by cloning and Sanger sequencing.**

PCR was carried out on DNA harvested from the heart tissue of dKO mice treated via intraperitoneal (IP) or intravenous (IV) injection of CRISPR AAVs with primers specific to *Dmd* intron 22 and the CMV promoter of the SaCas9-AAV genome (**Figure 8D**). Amplicons obtained from one AAV-IP treated animal were cloned into shuttle vectors and seven single clones analyzed by Sanger sequencing. Fragments of the SaCas9-AAV vector backbone was detected in all clones tested. Uppercase nucleotides indicate insertions of random bases not present in the WT sequence.
